# Effects of One-week Intake of Different Edible Oils on the Urinary Proteome of Rats

**DOI:** 10.1101/2025.02.04.636392

**Authors:** Yan Su, Youhe Gao

## Abstract

**Objective:** To explore the effects of different edible oils on the rat body by analyzing the changes in the urinary proteome and post-translational modifications of rats after one-week intake of olive oil, butter, lard, hydrogenated vegetable oil, and rapeseed oil.

**Methods:** Six parallel experiments were set up using male Wistar rats. Group A served as the control group, while groups B - F were administered different edible oils. The daily intakes were calculated respectively according to the “2015 - 2020 Dietary Guidelines for Americans” and the “Dietary Guidelines for Chinese Residents”. Urine samples were collected after one week and identified by label - free quantitative proteomics technology of high - performance liquid chromatography - tandem mass spectrometry (LC - MS/MS). Differentially expressed proteins and differential post - translational modifications in the urinary proteome were screened for functional analysis.

**Results:** After the intake of different types of edible oils, rats produced a large number of proteins and biological pathways related to metabolism, but there were few common differentially expressed proteins among different groups. In addition, the olive oil group and the butter group were enriched with many biological pathways related to the nervous system, and the rapeseed oil group produced more differentially expressed proteins and biological pathways related to immunity.

**Conclusion:** The urinary proteome of rats showed significant changes after one-week intake of edible oils, and the effects of various edible oils on the rat urinary proteome were different from each other. This effect is comprehensive and multi-dimensional at the level of the rat body. The changes in post-translational modifications of the proteome were relatively small.

## 1 Introduction

As an important component of the daily diet, edible oils are widely consumed worldwide. They provide essential fatty acids and energy for the human body and play a crucial role in maintaining normal physiological functions of the human body^[1]^. With the increasing attention to health, the nutritional values and potential health impacts of different edible oils have gradually become research hotspots.

Previous studies have shown that different types of edible oils have different physiological functions and health effects due to their unique fatty acid compositions and nutritional components^[2]^. For example, olive oil, the main source of fat in the Mediterranean diet, is rich in monounsaturated fatty acids and is considered to have beneficial effects such as reducing the risk of cardiovascular diseases^[3]^. Butter contains a relatively high amount of saturated fatty acids, and excessive intake may be associated with an increased risk of cardiovascular diseases^[4]^. Lard is an animal oil consumed frequently by people in Asian regions. Its fatty acid composition is relatively balanced, including 52.1% saturated fatty acids, 35.8% monounsaturated fatty acids, and 11.6% polyunsaturated fatty acids, and its impact on health is controversial^[5][6]^. Hydrogenated vegetable oil may have an adverse impact on cardiovascular health due to the presence of trans-fatty acids^[7]^. Rapeseed oil, as a vegetable oil, has a relatively balanced fatty acid ratio and contains nutritional components such as antioxidants^[8]^.

Given the significant differences in fatty acid compositions and nutritional characteristics among different edible oils, as well as their common consumption in daily diets, this study selected several representative edible oils, including olive oil, butter, lard, hydrogenated vegetable oil, and rapeseed oil, for research. To avoid the interference of short-term growth and development of rats on the experimental results, we chose to conduct parallel comparisons between different experimental groups and the control group. By comparing the differences in the urinary proteomes of rats, we preliminarily explored the possible impacts of short-term consumption of different edible oils on the metabolic processes in rats, providing certain theoretical references and experimental bases for a deeper understanding of their potential impacts on human health.

## 2 Introduction

### 2.1 Experimental Animals and Model Establishment

Thirty 7-week-old male Wistar rats, with a body weight of approximately 200 g, were purchased from Beijing Vital River Laboratory Animal Technology Co., Ltd. All rats were raised in a standard environment (room temperature: (22 ± 1) °C, humidity: 65%-70%). The experiment started after the rats were raised in the new environment for three days. All experimental operations were reviewed and approved by the Ethics Committee of the College of Life Sciences, Beijing Normal University, with the approval number CLS-AWEC-B-2022-003.

In this experiment, group A was set as the control group, without additional intake of edible oil. The other groups were fed with different types of edible oils, namely olive oil (group B), butter (group C), lard (group D), hydrogenated vegetable oil (group E), and rapeseed oil (group F), with 5 rats in each group. According to the “2015-2020 Dietary Guidelines for Americans” and the “Dietary Guidelines for Chinese Residents”, the ratio of the daily intake of edible oil to body weight for adults was calculated. Then, based on the equivalent dose ratio table for humans and animals converted according to body surface area, the dosage for rats was approximately 6.25 times that for humans. The daily intake of edible oil for rats was calculated as shown in Table 1.

**Table 1.** Daily Intake of Edible Oil for Rats.

|  | Olive Oil | Butter | Lard | Hydrogenated<br>Vegetable Oil | Rapeseed<br>Oil |
| --- | --- | --- | --- | --- | --- |
| Ratio of adult edible oil intake to<br>body weight / ((g/Kg)/day) | 0.5 | 0.44 | 0.45 | 0.45 | 0.45 |
| Rat edible oil intake / (g/day) | 0.625 | 0.55 | 0.56 | 0.56 | 0.56 |

After the adaptive feeding period ended, at 17:00 every day, rats in different groups were fed with the corresponding edible oils for 7 consecutive days. On the 7th day, after the rats consumed the edible oils, they were placed in metabolic cages to metabolize. After 12 hours overnight, urine was collected. Finally, the collected urine samples were centrifuged at 3000 r/min for 30 minutes and then stored in a-80°C refrigerator.

### 2.2 Processing of Urine Samples

Urine Protein Extraction Procedure: First, thaw the urine samples, then centrifuge them at 12000×g for 30 minutes at 4°C. Transfer the supernatant to a new EP tube. After that, add four-fold volume of absolute ethanol, mix well, and place it in a-20°C refrigerator for one-day precipitation. The next day, centrifuge the mixture at 12000×g for 30 minutes at 4°C, discard the supernatant, resuspend the protein precipitate in lysis buffer (containing 8 mol/L urea, 2 mol/L thiourea, 25 mmol/L dithiothreitol, 50 mmol/L), centrifuge again at 12000×g for 30 minutes at 4°C, and retain the supernatant. Finally, determine the protein concentration using the Bradford method.

Protease Digestion Procedure: Take 100 μg of urine protein sample and put it into an EP tube. Add 25 mmol/L NH□HCO□ solution to make the total volume 200 μL as a standby sample. Then add dithiothreitol solution (Dithiothreitol, DTT, Sigma) to the tube until its final concentration is 20 mM. Place it in a 95°C metal bath for 10 minutes of heating, then cool it to room temperature. Next, add iodoacetamide (Iodoacetamide, IAA, Sigma) to a concentration of 50 mM, and let it stand in the dark at room temperature for 40 minutes. Subsequently, wash the membrane of a 10-kDa ultrafiltration tube (Pall, Port Washington, NY, USA): First, add 200 μL of UA solution (8 mol/L urea, 0.1 mol/L Tris-HCl, pH 8.5), and centrifuge at 14000×g for 10 minutes at 18°C, repeat twice. Add the prepared sample to the ultrafiltration tube and centrifuge at 14000×g for 40 minutes at 18°C. Then add 200 μL of UA solution and centrifuge at 14000×g for 30 minutes at 18°C, repeat twice. Subsequently, add 25 mmol/L NH□HCO□ solution and centrifuge at 14000×g for 30 minutes at 18°C, repeat twice. Finally, add trypsin (Trypsin Gold, Promega, Fitchburg, WI, USA) at a ratio of trypsin:protein = 1:50, and incubate overnight at 37°C. After overnight incubation, centrifuge to collect the digested filtrate, desalt it using an Oasis HLB solid-phase extraction column, and vacuum-dry it to obtain freeze-dried peptides, which are stored at-80°C.

### 2.3 LC-MS/MS Tandem Mass Spectrometry Analysis

Place the peptides in 0.1% formic acid water for re-dissolution, and adjust the peptide concentration to 0.5 μg/μL through this operation. Subsequently, the peptides are separated by a Thermo ESAY-Nlc1200 liquid-phase system. The parameter settings of this liquid-phase system are as follows: The elution time is controlled at 90 minutes, and the elution gradient is specifically composed of phase A (0.1% formic acid) and phase B (80% acetonitrile). Finally, use an Orbitrap Fusion Lumos Tribird mass spectrometer to collect data-dependent mass spectrometry data for the separated peptides multiple times. The number of repeated collections is three times, and the data-independent acquisition mode is adopted.

### 2.4 Database Search

Use the Spectronaut Pulsar (Biognosys AG, Switzerland) software to search the database for the raw files of the mass spectrometer acquisition results, and compare them with the SwissProt Human database. Calculate the peptide abundance by adding the peak areas of the respective fragment ions in MS2. The protein intensity is calculated by adding the abundances of the respective peptides to calculate the protein abundance.

### 2.5 Open-pFind Unrestricted Modification Search

Use the pFind Studio software (version 3.2.1, Institute of Computing Technology, Chinese Academy of Sciences) to perform an unrestricted modification search on the three technical replicates of each sample. Use the default parameter settings during the search. The database used is the Rattus norvegicus database downloaded from UniProt, which has been updated to September 2024. The instrument type is set as HCD-FTMS, the selected enzyme is trypsin with full enzyme specificity, allowing a maximum of 2 missed cleavage sites. The precursor mass tolerance is set to ±20 ppm, the fragment mass tolerance is also set to ±20 ppm, and the open-search mode is selected. The screening condition clearly stipulates that the false discovery rate (FDR) at the peptide level should be less than 1%.

### 2.6 Bioinformatics Analysis of Protein Data

Each sample undergoes three technical replicates. The obtained data are averaged and used for statistical analysis. In this experiment, group comparisons were carried out between the edible oil groups (groups B, C, D, E, F) and the control group (group A) to screen for differentially expressed proteins. The screening criteria for differentially expressed proteins are: fold-change (FC) between groups ≥ 1.5 or ≤ 0.67, and the P-value of two-tailed paired t-test analysis < 0.05. The names and functions of the screened differentially expressed proteins are queried through the Uniprot website (https://www.uniprot.org), and biological function enrichment analysis is performed through the DAVID database (https://david.ncifcrf.gov). Also, retrieve reported literature in the Pubmed database (https://pubmed.ncbi.nlm.nih.gov) to conduct functional analysis of the differentially expressed proteins.

### 2.7 Bioinformatics Analysis of Protein Post-translational Modifications

Use Open-pFind to carry out an unrestricted modification search to obtain the post-translational modification PROTEIN file for each sample. Subsequently, download the Python script named pFind_protein_contrast_script on the GitHub platform (the website is https://github.com/daheitu/scripts_for_pFind3_protocol.io), and use this script to summarize the post-translational modification identification results (i.e., the PROTEIN files) of different samples. Similarly, group comparisons were carried out between the edible oil groups (groups B, C, D, E, F) and the control group (group A) to screen for differential modifications. The screening criteria for differential modifications are: FC ≥ 1.5 or ≤ 0.67, and the P-value of two-tailed paired t-test analysis < 0.05. Query the names and functions of the proteins where the screened differential modifications are located through the Uniprot website (https://www.uniprot.org).

## 3 Results and Discussion

### 3.1 Behavioral Observation of Rats

The body weights of rats were measured before and after the experiment (Figure 1a). By comparison, it was found that the body weight gain of rats in the experimental groups, except for group E, was higher than that of the control group. The food intake of rats during the experiment was counted (Figure 1b), and it was found that the food intake of rats in the experimental groups was higher than that of the control group. When observing the preference of rats for edible oils, it was found that rats in groups C and D consumed butter and lard more quickly, while rats in group E obviously rejected the hydrogenated vegetable oil they were fed.

**Figure 1.**
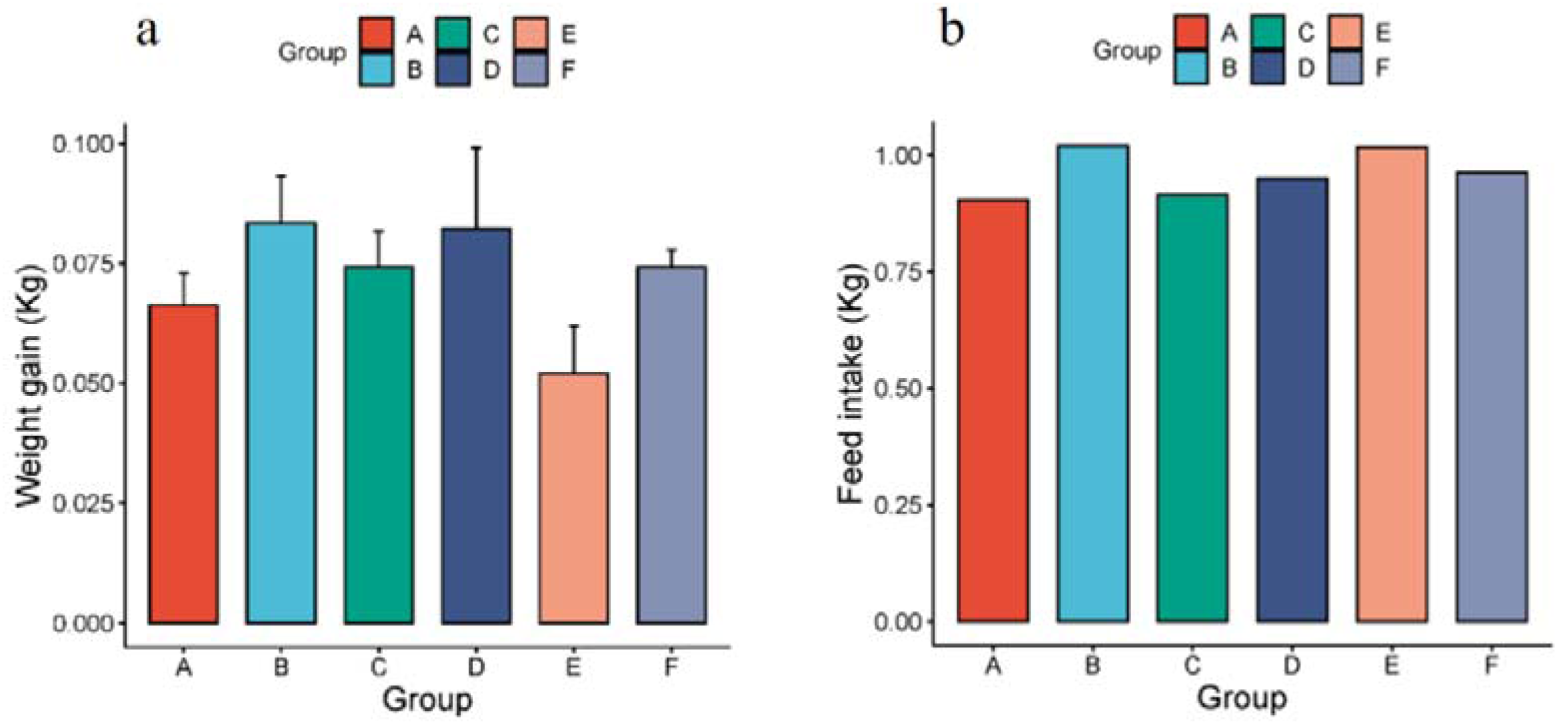
a. Body weight gain of rats before and after oil-feeding; b. Food intake of rats during oil-feeding.

### 3.2 Group Analysis of Urinary Protein Composition

#### 3.2.1 Random Grouping

The control group samples (n = 5) and the experimental group samples (n = 5) were randomly divided into two groups, with a total of 126 grouping types. Among all the random combination types, the average number of differentially expressed proteins for all random times was calculated according to the same screening criteria. The ratio of the average number of differentially expressed proteins to the number of differentially expressed proteins obtained under normal grouping is the proportion of randomly generated differentially expressed proteins, as listed in Table 2. These results indicate that the number of randomly generated differentially expressed proteins is relatively small, and the reliability of screening differentially expressed proteins is high. It shows that most of the differentially expressed proteins we obtained are not randomly generated but are the result of the impact of short-term intake of different edible oils.

**Table 2.** Proportion of Randomly Generated Differentially Expressed Proteins Obtained from Random Grouping.

| Control-Experimental Group | A-B | A-C | A-D | A-E | A-F |
| --- | --- | --- | --- | --- | --- |
| Proportion of Randomly Generated Proteins | 5.6% | 6.6% | 6.4% | 12.7% | 5.5% |

#### 3.2.2 Group Analysis of Urinary Protein Composition in the Olive Oil Group

##### 3.2.2.1 Differentially Expressed Proteins

The urinary proteins of the olive oil group (group B) and the control group (group A) were compared. The screening criteria for differentially expressed proteins were: fold-change FC ≥ 1.5 or ≤ 0.67, and two-tailed paired t-test P < 0.05. The results showed that, compared with the control group, a total of 94 differentially expressed proteins were identified in the olive oil group, among which 17 were down-regulated and 77 were up-regulated. The differentially expressed proteins were arranged in ascending order of FC, and retrieved through Uniprot. The detailed information is listed in Appendix Table 1.

##### 3.2.2.2 Functional Analysis of Differentially Expressed Proteins

The identified differentially expressed proteins were searched in the Pubmed database for relevant literature.

Among the differentially expressed proteins, the one with the largest down-regulation amplitude and a change from “present to absent” was Alpha-1,3-mannosyl-glycoprotein 4-beta-N-acetylglucosaminyltransferase A (FC = 0, P = 3.81E-02). Research shows that N-acetylglucosaminyltransferase IV is responsible for transferring uridine diphosphate-N-acetylglucosamine to specific glycoprotein structures and is involved in the glycosylation modification of tumor-related N-glycans. This process may be associated with the pathogenesis of diabetes^[9]^.

In addition, some proteins have been shown in existing literature to be related to olive oil, oxidative stress, or metabolism:

IQ motif containing GTPase activating protein 2 (FC = 0.094, P = 2.22E-02) plays an important role in regulating liver fuel storage, especially glycogen levels. In female mice lacking this protein, a significant reduction in glycogen accumulation was observed^[10]^.

Selenoprotein P (FC = 0.213, P = 1.71E-03), as the main selenium-transporting protein, plays a crucial role in maintaining selenium homeostasis in the body and the hierarchical structure of selenoprotein expression. Adequate selenium supply is essential for the normal development and function of the brain through SELENOP, and is crucial for male fertility, proper neurological function, and selenium metabolism^[11]^.

Heat shock protein HSP 90-beta (FC = 0.318, P = 8.62E-03) is abundantly expressed in cardiomyocytes. By maintaining the level of glutathione (a major redox mediator), it is involved in maintaining the redox homeostasis of the cardiovascular system^[12]^.

Glutathione peroxidase (FC = 3.773, P = 2.25E-03) is an important antioxidant enzyme that plays a key role in protecting cells from oxidative stress damage. Studies have found that ligstroside, a polyphenolic compound in olive oil, can increase the expression of Glutathione peroxidase, which may help improve the antioxidant capacity of cells and thus protect cells from damage caused by oxidative stress^[13]^.

Angiopoietin-like 2 (FC = 7.891, P = 1.18E-02) is an important angiogenic factor. In recent years, it has also been found to be involved in mediating the inflammatory process. Its expression is up-regulated in various inflammatory diseases and is directly related to the regulation of inflammatory-related signaling pathways^[14]^.

Leucine-rich repeat transmembrane protein FLRT2 (FC = 8.891, P = 5.15E-03) promotes lipid peroxidation by increasing the expression of acyl-CoA synthetase long-chain family member 4, thereby triggering ferroptosis and inhibiting the malignant phenotype of human bladder cancer cells. It has been identified as a tumor-suppressor gene^[15]^.

Oxidized purine nucleoside triphosphate hydrolase (FC = 10.262, P = 2.83E-03) hydrolyzes oxidized purine nucleoside triphosphates (such as 8-oxo-dGTP and 2-hydroxy-dATP) to monophosphates, thus preventing the mis-incorporation of these oxidized nucleotides during replication^[16]^.

##### 3.2.2.3 Enrichment Analysis of Biological Processes of Differentially Expressed Proteins

The DAVID database was used to perform enrichment analysis of biological processes (BP) on the 94 differentially expressed proteins identified by group analysis. The results showed that a total of 29 BP pathways were significantly enriched (P < 0.05), and the detailed information is presented in Figure 2.

**Figure 2.**
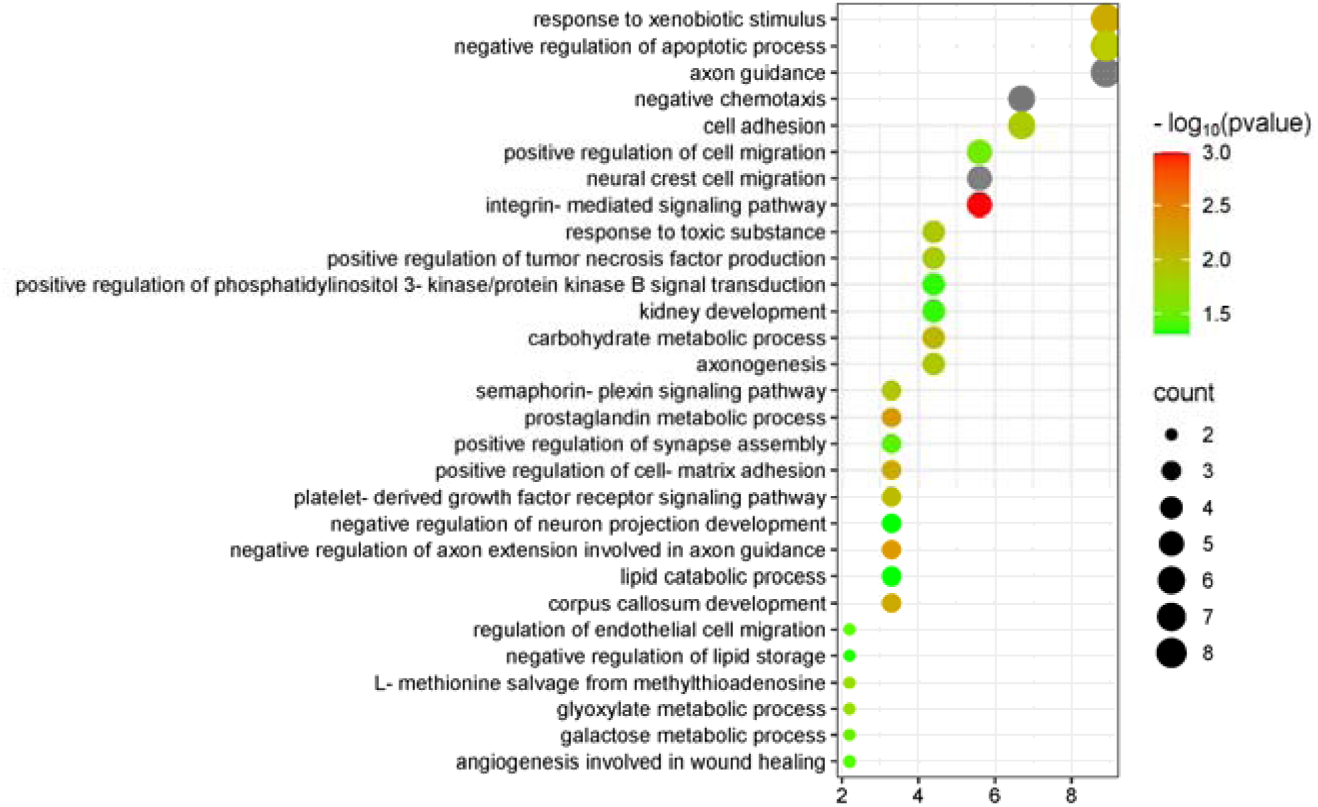
Biological Processes Enriched by Differentially Expressed Proteins between the Olive Oil Group and the Control Group

Many of the enriched pathways are related to the nervous system, including axon guidance, neural crest cell migration, negative regulation of axon extension involved in axon guidance, corpus callosum development, axogenesis, positive regulation of synapse assembly, and negative regulation of neuron projection development. Multiple studies have shown that extra-virgin olive oil has important neuroprotective activities, which are closely related to its specific phenolic components. Therefore, nervous-system-related pathways may be enriched^[17][18][19]^.

The pathway “regulation of endothelial cell migration” has also been mentioned in olive oil research: Hydroxytyrosol and secoiridoids are both components of olive oil, and they play a role in regulating proteins related to endothelial cell proliferation and migration and in regulating proteins related to heart failure in cardiac tissue^[20]^.

In addition, the negative regulation of lipid storage, lipid catabolic process, and negative regulation of lipid biosynthetic process are three pathways directly related to fat intake.

#### 3.2.3 Group Analysis of Urinary Protein Composition in the Butter Group

##### 3.2.3.1 Differentially Expressed Proteins

The urinary proteins of the butter group (group C) and the control group (group A) were compared. The screening criteria for differentially expressed proteins were: fold-change FC ≥ 1.5 or ≤ 0.67, and two-tailed paired t-test P < 0.05. The results showed that, compared with the control group, a total of 89 differentially expressed proteins were identified in the butter group, among which 15 were down-regulated and 74 were up-regulated. The differentially expressed proteins were arranged in ascending order of FC, and retrieved through Uniprot. The detailed information is listed in Appendix Table 2.

##### 3.2.3.2 Functional Analysis of Differentially Expressed Proteins

The identified differentially expressed proteins were searched in the Pubmed database for relevant literature. Some of the differentially expressed proteins in the butter group were the same as those in the olive oil group, such as Heat shock protein HSP 90-beta, Glutathione peroxidase, Oxidized purine nucleoside triphosphate hydrolase, etc.

Among the other down-regulated differentially expressed proteins in the butter group: C-C motif chemokine 2 (FC = 0.190, P = 2.18E-02) plays a major role in the pathogenesis of cardiovascular diseases. Its down-regulation is considered one of the pleiotropic properties of statins^[21]^. Neuronal membrane glycoprotein M6-a (FC = 0.277, P = 2.56E-02) plays an important role in activating the Src/MAPK/ERK, PKC, and PI3K/AKT, Rufy3-Rap2-STEF/Tiam2 signaling pathways, promoting neurite outgrowth and neuronal polarization respectively, and is also involved in the formation and maturation of dendritic spines^[22]^. Selenoprotein P (FC = 0.340, P = 2.72E-04), as the main selenium-transporting protein, maintains an adequate selenium supply in the brain, which is crucial for the normal development and function of the brain. It may also be related to the pathological processes of the central nervous system and has antioxidant activity^[23]^.

Among the up-regulated proteins: Metalloproteinase inhibitor 3 (FC = 306.177, P = 1.05E-02) had the largest up-regulation amplitude. Studies have found that overexpression of TIMP3 inhibited pathways related to metabolic inflammation and stress, including the activation of Jun NH2-terminal kinase and p38 kinase, and reduced the activation of oxidative stress signals, which were related to lipid peroxidation, protein carbonylation, and nitration^[24]^. An increase in the activity of Prenylcysteine oxidase 1 (FC = 17.347, P = 4.97E-02) may lead to an increase in hydrogen peroxide, thus increasing the oxidative burden during the propagation of low-density lipoproteins. Therefore, this protein can serve as a potential drug target and a new biomarker for cardiovascular diseases^[25]^. Cellular repressor of E1A-stimulated genes 1 (FC = 6.517, P = 3.59E-02) may regulate the homeostasis of vascular wall cells and inhibit the inflammation of vascular tissue cells and macrophages, and has a potential protective effect against inflammation^[26]^. Palmitoyl-protein thioesterase 1 (FC = 3.266, P = 1.42E-02) was up-regulated in rats on a high-fat diet. The increase in its expression may be harmful to the function of Sertoli cells during spermatogenesis^[27]^.

##### 3.2.3.3 Enrichment Analysis of Biological Processes of Differentially Expressed Proteins

The DAVID database was used to perform enrichment analysis of biological processes (BP) on the 89 differentially expressed proteins identified by group analysis. The results showed that a total of 40 BP pathways were significantly enriched (P < 0.05), and the detailed information is presented in Figure 3.

**Figure 3.**
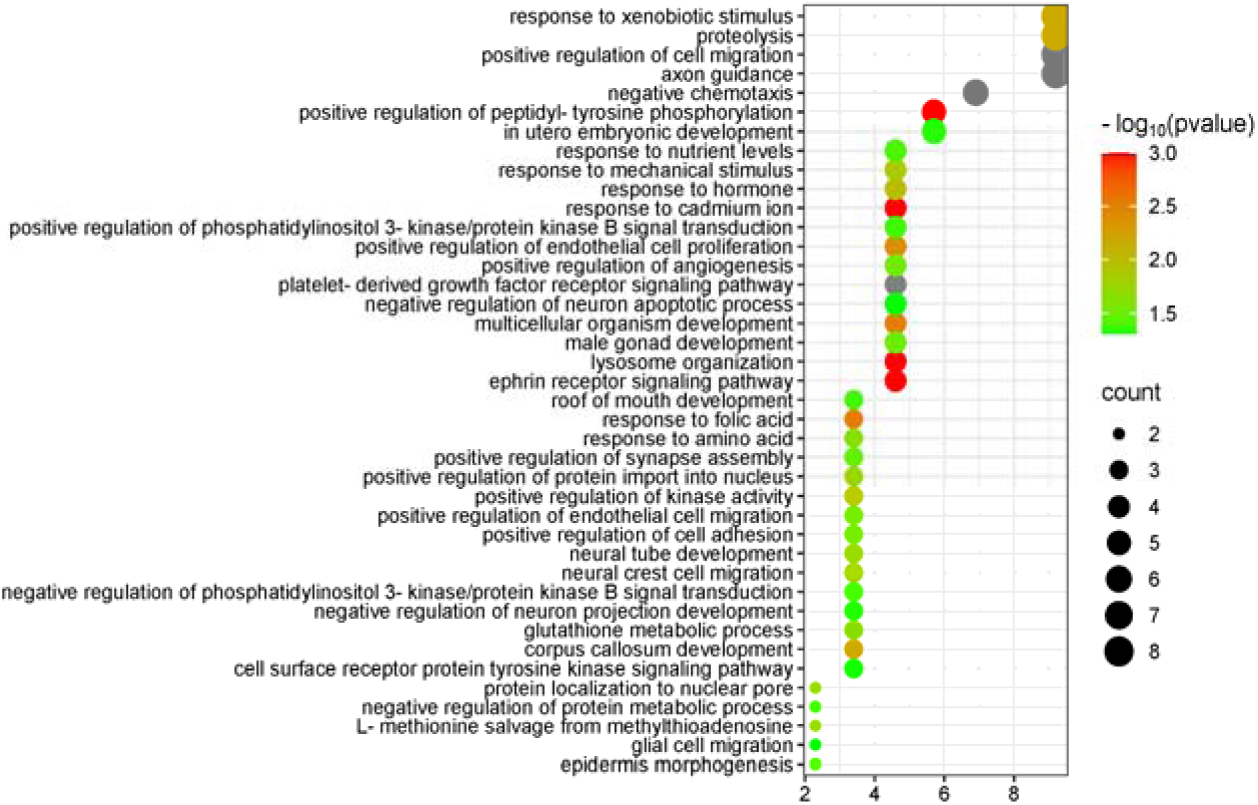
Biological Processes Enriched by Differentially Expressed Proteins between the Butter Group and the Control Group

Similar to the olive oil group, the butter group was also enriched with many pathways related to the nervous system, including axon guidance, corpus callosum development, neural crest cell migration, etc. In addition, the butter group was enriched with some metabolism-related pathways, such as proteolysis, glutathione metabolic process, response to nutrient levels, and negative regulation of protein metabolic process.

#### 3.2.4 Group Analysis of Urinary Protein Composition in the Lard Group

##### 3.2.4.1 Differentially Expressed Proteins

The urinary proteins of the lard group (group D) and the control group (group A) were compared. The screening criteria for differentially expressed proteins were: fold-change FC ≥ 1.5 or ≤ 0.67, and two-tailed paired t-test P < 0.05. The results showed that, compared with the control group, a total of 104 differentially expressed proteins were identified in the lard group, among which 25 were down-regulated and 79 were up-regulated. The differentially expressed proteins were arranged in ascending order of FC, and retrieved through Uniprot. The detailed information is listed in Appendix Table 3.

##### 3.2.4.2 Functional Analysis of Differentially Expressed Proteins

The identified differentially expressed proteins were searched in the Pubmed database for relevant literature.

Among the down-regulated differentially expressed proteins in the lard group: Mitogen-activated protein kinase 3 (FC = 0, P = 2.67E-02) may play an important role in the functional regulation of neurons. Its regulation of the mGlu5 receptor is involved in various neurobiological processes, such as synaptic plasticity, learning, and memory ^[28]^. Protein S100-A1 (FC = 0.111, P = 2.99E-02) undergoes a large conformational change when binding to calcium to interact with numerous protein targets, including proteins involved in calcium signaling, neurotransmitter release (synapsins I and II), and the cytoskeleton^[29]^. Dickkopf WNT signaling pathway inhibitor 3 (FC = 0.120, P = 4.74E-02) is significantly down-expressed in the brain tissues of Alzheimer’s disease patients and transgenic mouse models of Alzheimer’s disease^[30]^.

Among the up-regulated differentially expressed proteins: The protein with the largest up-regulation amplitude was Metalloproteinase inhibitor 4 (FC = 85.055, P = 3.60E-02), which belongs to the family of extracellular matrix metalloproteinase inhibitors and is over-expressed in various cancers. Currently, there is no research on its association with lard intake^[31]^. Retinol-binding protein 1 (FC = 38.811, P = 3.04E-02), as a chaperone protein, regulates the uptake, subsequent esterification, and bioavailability of retinol and can deliver vitamin A to cells by interacting with cell-membrane receptors^[32][33]^. Hyaluronan and proteoglycan link protein 1 (FC = 13.005, P = 5.51E-03) inhibits the NLRP3 inflammasome by stimulating the Nrf2/ARE pathway, thereby suppressing neuroinflammation, enhancing motor neuron survival, and improving neurological functional recovery after spinal cord injury^[34]^. Prominin-2 (FC = 8.125, P = 4.53E-02) is induced by ferroptosis stimulation and can play a role in resisting ferroptosis^[35]^. Protein disulfide-isomerase A6 (FC = 8.080, P = 4.80E-02) is related to spinal cord injury repair and has also been found to promote the repair of damaged neurons^[36]^. 2-amino-3-carboxymuconate-6-semialdehyde decarboxylase (FC = 7.322, P = 1.31E-03) plays a key role in tryptophan catabolism and is an attractive therapeutic target for treating diseases associated with elevated levels of tryptophan metabolites^[37]^.

##### 3.2.4.3 Enrichment Analysis of Biological Processes of Differentially Expressed Proteins

The DAVID database was used to perform enrichment analysis of biological processes (BP) on the 104 differentially expressed proteins identified by group analysis. The results showed that a total of 32 BP pathways were significantly enriched (P < 0.05), and the detailed information is presented in Figure 4.

**Figure 4.**
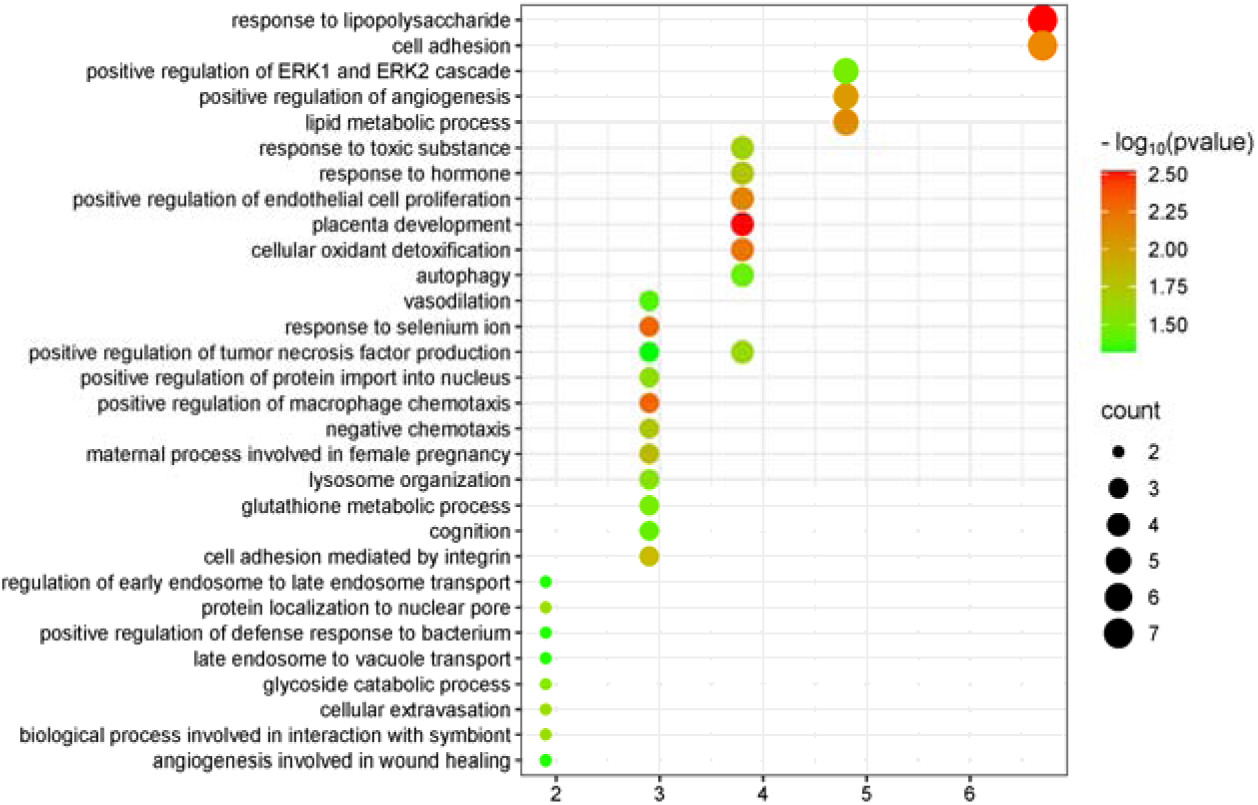
Biological Processes Enriched by Differentially Expressed Proteins between the Lard Group and the Control Group

Among the enriched pathways, there were some metabolism-related pathways, including the response to lipopolysaccharide, lipid metabolic process, glycoside catabolic process, and glutathione metabolic process. The nervous-system-related pathways that appeared frequently in the olive oil group and the butter group did not appear.

#### 3.2.5 Group Analysis of Urinary Protein Composition in the Hydrogenated Vegetable Oil Group

##### 3.2.5.1 Differentially Expressed Proteins

The urinary proteins of the hydrogenated vegetable oil group (group E) and the control group (group A) were compared. The screening criteria for differentially expressed proteins were: fold-change FC ≥ 1.5 or ≤ 0.67, and two-tailed paired t-test P < 0.05. The results showed that, compared with the control group, a total of 63 differentially expressed proteins were identified in the hydrogenated vegetable oil group, among which 34 were down-regulated and 29 were up-regulated. The differentially expressed proteins were arranged in ascending order of FC, and retrieved through Uniprot. The detailed information is listed in Appendix Table 4.

##### 3.2.5.2 Functional Analysis of Differentially Expressed Proteins

The identified differentially expressed proteins were searched in the Pubmed database for relevant literature.

Among the down-regulated differentially expressed proteins in the hydrogenated vegetable oil group: Disabled homolog 2 (FC = 0, P = 4.72E-02) plays a crucial role in controlling macrophage phenotypic polarization and adipose tissue inflammation. Its deficiency leads to exacerbated adipose tissue inflammation and insulin resistance^[38]^. Muscles with knockout of Calmodulin-like protein 3 (FC = 0.011, P = 4.55E-02) gene show weakened calcium release, reduced calmodulin kinase signaling, and impaired muscle adaptation to exercise^[39]^. Integrin alpha-1 (FC = 0.036, P = 2.67E-02) can promote hepatic insulin action and lipid accumulation simultaneously under high-fat diet challenge, and plays a role in liver metabolism. Its deficiency can alter fatty acid metabolism in high-fat diet mice and improve fatty liver conditions^[40]^. Alpha-2-HS-glycoprotein (FC = 0.116, P = 4.48E-02) is a multifunctional plasma glycoprotein mainly synthesized in the liver. It is considered an important part of various normal and pathological processes, including bone metabolism regulation, vascular calcification, insulin resistance, and protease activity control. During obesity and related complications such as type 2 diabetes, metabolic syndrome, and non-alcoholic fatty liver disease, the circulating level of Alpha-2-HS-glycoprotein is significantly increased^[41]^. Eppin (FC = 0.192, P = 2.36E-02) is a biomarker of androgen action in human semen. Research has found that it is closely related to semen quality, sperm motility, etc. Down-regulation of Eppin expression significantly reduces sperm motility^[42]^.

Among the up-regulated differentially expressed proteins: Phosphoserine phosphatase (FC = 14.636, P = 4.44E-02) is the protein with the largest up-regulation amplitude. No literature research has shown its correlation with metabolic activities. Catechol O-methyltransferase (FC = 3.841, P = 2.40E-02) plays a key role in metabolizing dopamine. Common functional polymorphisms in the gene of this protein affect cognitive functions related to the prefrontal cortex, sleep-wake regulation, and may affect sleep pathology^[43]^.

##### 3.2.5.3 Enrichment Analysis of Biological Processes of Differentially Expressed Proteins

The DAVID database was used to perform enrichment analysis of biological processes (BP) on the 63 differentially expressed proteins identified by group analysis. The results showed that a total of 16 BP pathways were significantly enriched (P < 0.05), and the detailed information is presented in Figure 5.

**Figure 5.**
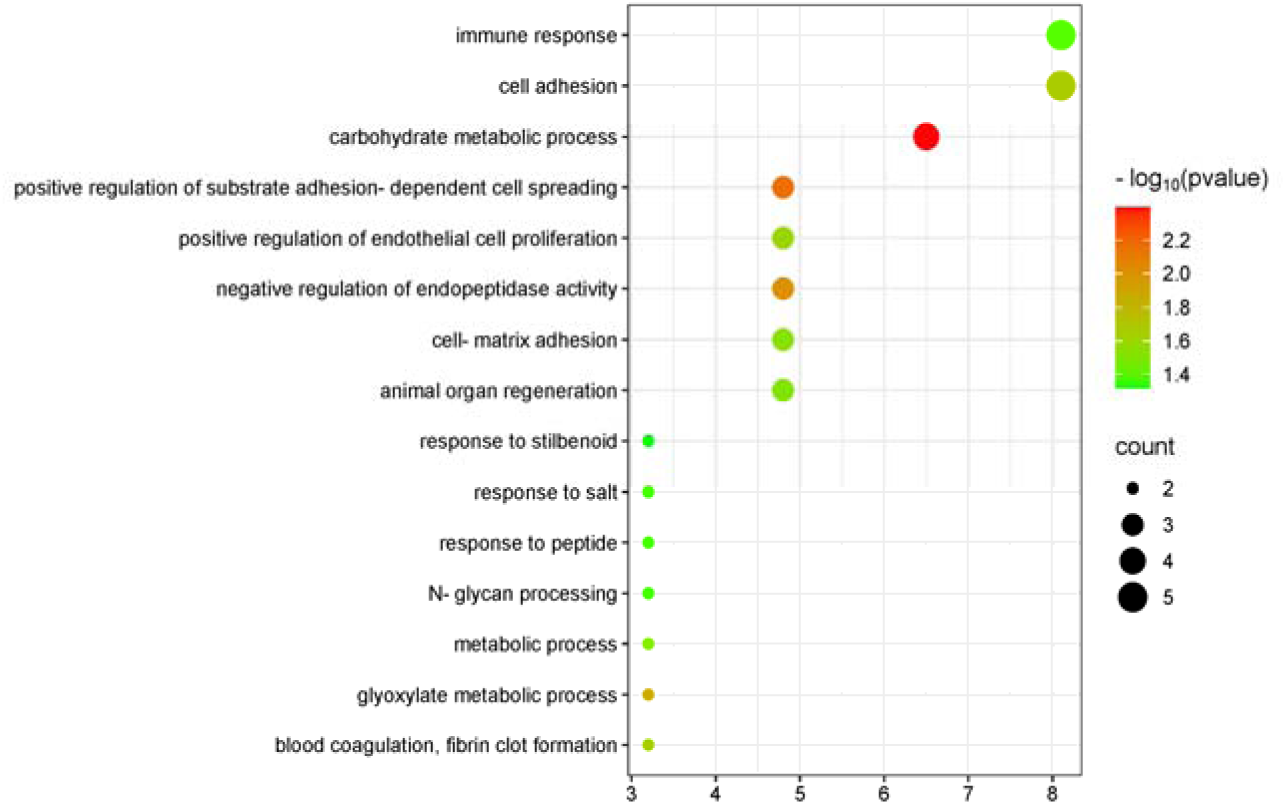
Biological Processes Enriched by Differentially Expressed Proteins between the Hydrogenated Vegetable Oil Group and the Control Group

Among the enriched pathways, there were some metabolism-related pathways, including carbohydrate metabolic process, glyoxylate metabolic process, and metabolic process. Also, the nervous-system-related pathways that appeared frequently in the olive oil group and the butter group were not present.

#### 3.2.6 Group Analysis of Urinary Protein Composition in the Hydrogenated Vegetable Oil Group

##### 3.2.6.1 Differentially Expressed Proteins

The urinary proteins of the rapeseed oil group (group F) and the control group (group A) were compared. The screening criteria for differentially expressed proteins were: fold-change FC ≥ 1.5 or ≤ 0.67, and two-tailed paired t-test P < 0.05. The results showed that, compared with the control group, a total of 105 differentially expressed proteins were identified in the rapeseed oil group, among which 53 were down-regulated and 52 were up-regulated. The differentially expressed proteins were arranged in ascending order of FC, and retrieved through Uniprot. The detailed information is listed in Appendix Table 5.

##### 3.2.6.2 Functional Analysis of Differentially Expressed Proteins

The identified differentially expressed proteins were searched in the Pubmed database for relevant literature.

Among the down-regulated differentially expressed proteins in the rapeseed oil group, a total of 20 differentially expressed proteins had the largest down-regulation amplitude with a change from “present to absent”. Existing literature has proven that multiple of these proteins are related to immune-inflammation, oxidative stress, etc. Thrombospondin 4 (FC = 0, P = 4.46E-02) is a stress-inducible secreted glycoprotein that plays a crucial role in tissue injury and healing. It can activate the adaptive endoplasmic reticulum stress response and enhance sarcolemma stability in the heart and skeletal muscles. Defective flux of Thrombospondin 4 through the secretory pathway impairs the stability of the cardiomyocyte membrane and leads to cardiomyopathy^[44]^. Interleukin 10 receptor subunit beta (FC = 0, P = 2.63E-02) is one of the components of the Interleukin 10 receptor, which is common to all members of the IL-10 cytokine family. Interleukin-10 is a type 2 T-helper cell cytokine with extensive anti-inflammatory effects^[45][46]^. Ring finger protein 150 (FC = 0, P = 2.83E-02) may be involved in regulating the antioxidant stress signaling pathway in neurons, enhancing the resistance of neurons to oxidative stress, and thus reducing neuronal damage and death^[47]^. Interleukin 17 receptor A (FC = 0, P = 2.50E-02) is the receptor for interleukin-17A, which is a pro-inflammatory cytokine related to the rapid malignant progression and treatment resistance of colorectal cancer^[48]^. Endothelial protein C receptor (FC = 0, P = 3.81E-02) plays a positive role in normal homeostasis, anticoagulant pathways, inflammation, and cell stemness, and is considered a potential effector or mediator of inflammatory diseases^[49]^. The only known functional ligand of Cell surface glycoprotein CD200 receptor 1 (CD200R) (FC = 0, P = 4.12E-02) is CD200, and their interaction leads to the activation of anti-inflammatory signals in CD200R-expressing cells. When this interaction becomes insufficient due to aging or disease, chronic inflammation occurs^[50]^. In addition, Protein O-linked-mannose beta-1,2-N-acetylglucosaminyltransferase 1 (POMGnT1) (FC = 0, P = 3.26E-02) is mainly expressed in neurons and also to a certain extent in glial cells. The expression of POMGnT1 decreases in mouse and cell models of Alzheimer’s disease^[51]^.

Among the up-regulated differentially expressed proteins, Serine peptidase inhibitor Kunitz type 3 (FC = 43.172, P = 4.78E-02) had the largest fold-change. It is involved in processes such as blood coagulation and fibrinolysis, tumor immunity, inflammation regulation, and resistance to bacterial and fungal infections^[52]^. CCN family member 1 (FC = 9.008, P = 3.96E-02) is an extracellular matrix protein that has a potential role in wound healing, accelerating re-epithelialization by promoting keratinocyte migration and proliferation^[53]^.

##### 3.2.6.3 Enrichment Analysis of Biological Processes of Differentially Expressed Proteins

The DAVID database was used to perform enrichment analysis of biological processes (BP) on the 105 differentially expressed proteins identified by group analysis. The results showed that a total of 51 BP pathways were significantly enriched (P < 0.05), and the detailed information is presented in Figure 6.

**Figure 6.**
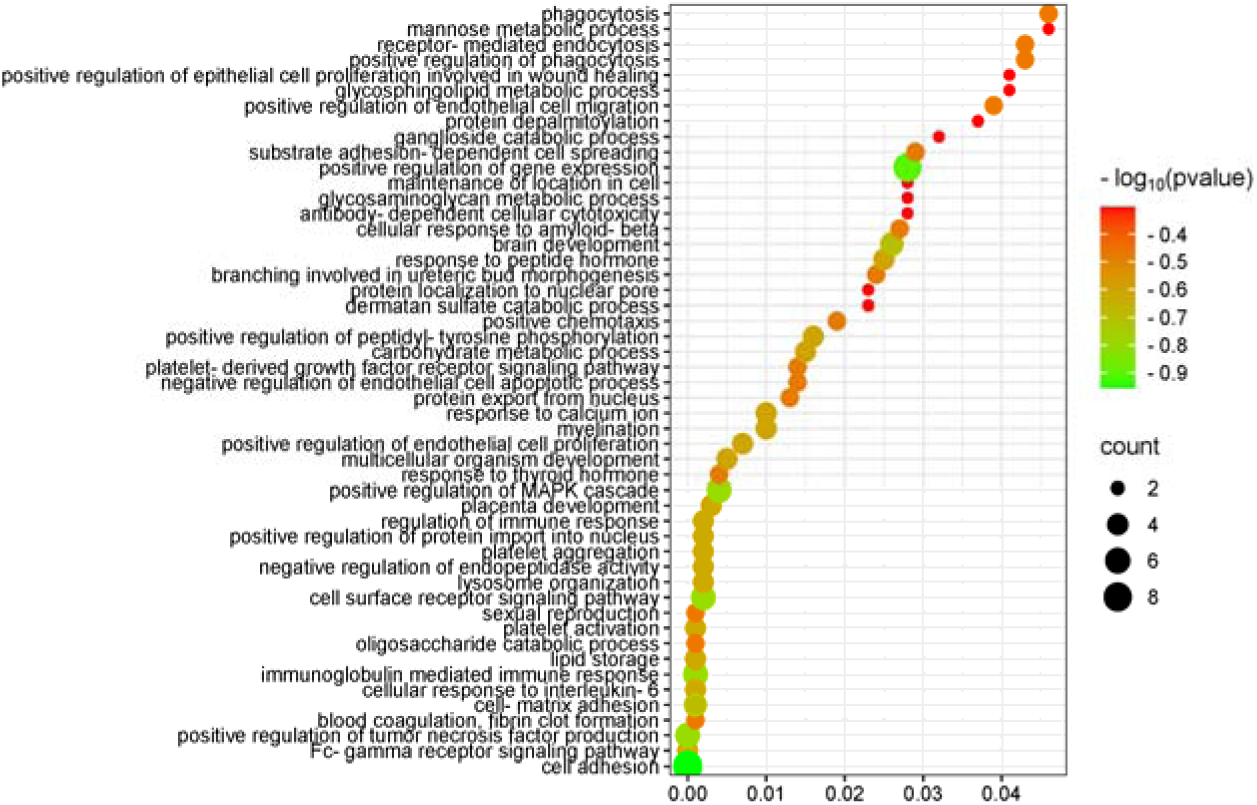
Biological Processes Enriched by Differentially Expressed Proteins between the Rapeseed Oil Group and the Control Group

**Figure 7.**
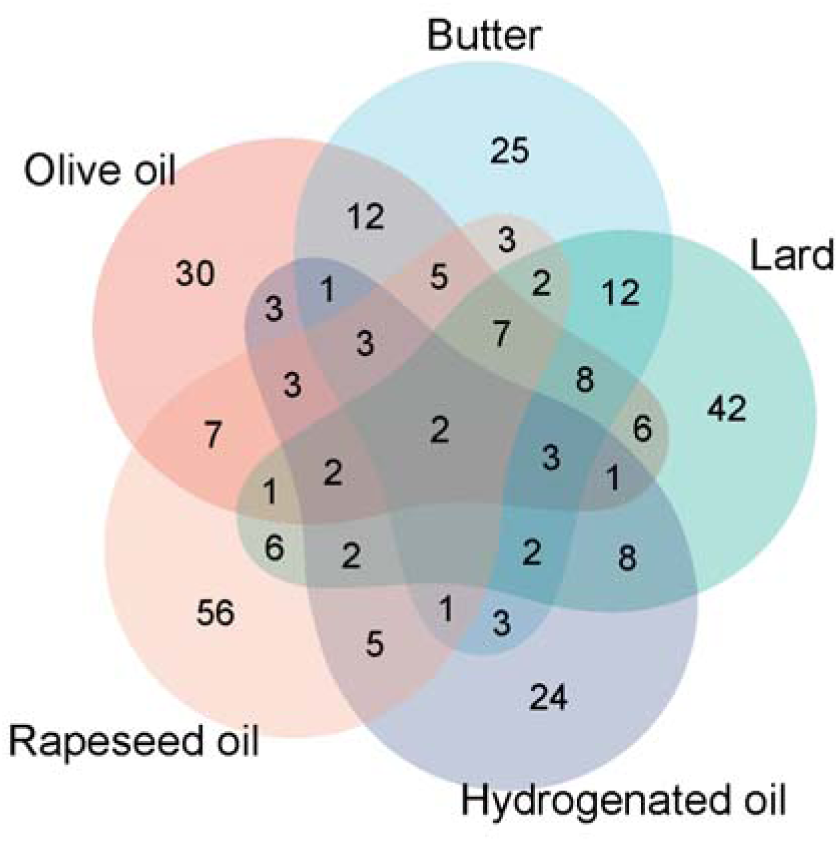
Common Differentially Expressed Proteins in the Urinary Proteomes of the Olive Oil Group, Butter Group, Lard Group, Hydrogenated Vegetable Oil Group, and Rapeseed Oil Group

Compared with other edible oil groups, the pathways enriched in the rapeseed oil group contained a large number of immune-related pathways, including the Fc-γ receptor signaling pathway, immunoglobulin-mediated immune response, cellular response to interleukin 6, regulation of the immune response, positive regulation of endothelial cell proliferation, negative regulation of the endothelial cell apoptotic process, antibody-dependent cell-mediated cytotoxicity, and positive regulation of endothelial cell migration. Studies have found that rapeseed oil shows certain positive effects in supporting the acquired immune capacity of weaned mice, especially in promoting the antibody response^[54]^.

In addition, many metabolism-related pathways were also enriched, such as lipid storage, oligosaccharide catabolic process, carbohydrate metabolic process, etc. It is worth noting the pathway “cellular response to beta-amyloid protein”. Amyloid-beta aggregates in the brain play a central role in the pathogenesis of Alzheimer’s disease, and research has shown that consumption of vegetable oil can be used as a preventive or adjuvant strategy to slow down or prevent the progression of neurodegenerative diseases^[55][56]^.

#### 3.2.7 Common Differentially Expressed Proteins in the Group Analysis of Different Edible Oils

The common differentially expressed proteins in the five-group experiments were counted. It was found that there were few common differentially expressed proteins, indicating that different edible oils have different impacts on the body. However, two proteins, Amyloid P component serum and 5’-3’ exonuclease PLD3, were up-regulated in the urine of all experimental groups. Amyloid P component serum is believed to potentially promote the progression of neurodegenerative diseases, including Alzheimer’s disease, by binding to β-amyloid protein to increase its stability and inducing neuronal apoptosis^[57]^. 5’-3’ exonuclease PLD3 is highly expressed in brain neurons. Its level is down-regulated in the brains of Alzheimer’s disease patients and is negatively correlated with the levels of amyloid-precursor protein and amyloid-β protein. PLD3 may be involved in the pathogenesis of Alzheimer’s disease through the processing of amyloid-precursor protein^[58]^.

### 3.3 Group Analysis of Urinary Post-translational Modifications

#### 3.3.1 Random Grouping

The control group samples (n = 5) and the experimental group samples (n = 5) were randomly divided into two groups, with a total of 126 grouping types. Among all the random combination types, the average number of post-translational modifications for all random times was calculated according to the same screening criteria. The ratio of the average number of post-translational modifications to the number of post-translational modifications obtained under normal grouping is the proportion of randomly generated post-translational modifications, as listed in Table 3. These results indicate that the proportion of randomly generated post-translational modifications is relatively large, suggesting that short-term intake of different edible oils has a relatively small impact on post-translational modifications in the rat body.

**Table 3.** Proportion of Randomly Generated Differential Modifications Obtained from Random Grouping.

| Control-Experimental Group | A-B | A-C | A-D | A-E | A-F |
| --- | --- | --- | --- | --- | --- |
| Proportion of Randomly Generated Proteins | 72.1% | 72.6% | 48.5% | 83.0% | 20.4% |

#### 3.3.2 Group Analysis of Post-translational Modifications in the Olive Oil Group

The post-translational modifications of the olive oil group and the control group were compared. The screening criteria for differential modifications were: FC≥1.5 or ≤0.67, and two-tailed paired t-test P < 0.05. The results showed that, compared with the control group, a total of 41 differential modifications were identified in the olive oil group, among which 17 were down-regulated and 24 were up-regulated. The differentially modified proteins were arranged in ascending order of FC, and the detailed information is listed in Appendix Table 6.

Among the proteins with differential modifications showing an up- or down-regulation of more than 10-fold: P97574, namely Stanniocalcin-1 (FC = 0, P = 4.59E-02), may be involved in the digestion and absorption in the gastrointestinal tract and kidneys of those parts with active digestion and absorption functions ^[59]^.

#### 3.3.3 Group Analysis of Post-translational Modifications in the Butter Group

The post-translational modifications of the butter group and the control group were compared. The screening criteria for differential modifications were: FC≥1.5 or ≤0.67, and two-tailed paired t-test P < 0.05. The results showed that, compared with the control group, a total of 49 differential modifications were identified in the butter group, among which 4 were down-regulated and 45 were up-regulated. The differentially modified proteins were arranged in ascending order of FC, and the detailed information is listed in Appendix Table 7.

Among the proteins with differential modifications showing an up- or down-regulation of more than 10-fold: P02770, namely Albumin (FC = 37, P = 1.87E-02), plays a crucial role in transporting various endogenous and exogenous molecules and maintaining the colloid osmotic pressure of blood. It has many enzymatic activities, and its free thiol groups determine that this protein can participate in redox reactions^[60]^. Q5FVF9, namely Biotinidase (FC = 24, P = 4.78E-02), is an enzyme that helps the body reuse and recycle biotin in food^[61]^.

#### 3.3.4 Group Analysis of Post-translational Modifications in the Lard Group

The post-translational modifications of the lard group and the control group were compared. The screening criteria for differential modifications were: FC≥1.5 or ≤0.67, and two-tailed paired t-test P < 0.05. The results showed that, compared with the control group, a total of 99 differential modifications were identified in the lard group, among which 23 were down-regulated and 76 were up-regulated. The differentially modified proteins were arranged in ascending order of FC, and the detailed information is listed in Appendix Table 8.

Among the proteins with differential modifications showing an up- or down-regulation of more than 10-fold: P07151, namely Beta-2-microglobulin (FC = 15, P = 8.64E-03), may play a role in the oxidative stress state of the elderly and can be used as a new biomarker of oxidative stress^[63]^. P11232, namely Thioredoxin (FC = 0.091, P = 3.41E-02), plays an important role in maintaining the redox state of cells and regulating redox signal transduction. Together with glutathione, it plays a central role in combating oxidative stress^[64]^.

#### 3.3.5 Group Analysis of Post-translational Modifications in the Hydrogenated Vegetable Oil Group

The post-translational modifications of the hydrogenated vegetable oil group were compared with those of the control group. The screening criteria for differential modifications were set as FC ≥ 1.5 or ≤ 0.67, and a two-tailed paired t-test with P < 0.05. The results indicated that, compared to the control group, a total of 50 differential modifications were identified in the hydrogenated vegetable oil group, among which 38 were down-regulated and 12 were up-regulated. The differentially modified proteins were arranged in ascending order of FC, and the detailed information is presented in Appendix Table 9.

Among the proteins with differential modifications showing an up- or down-regulation of more than 10-fold, P27590, namely Uromodulin (FC = 0.053, P = 4.91E-02), has multiple functions in kidney physiology, renal tubular transport, and mineral metabolism. A higher serum concentration of Uromodulin is associated with higher renal function and preserved renal reserve. It is also related to a lower risk of cardiovascular diseases and diabetes^[65]^.

#### 3.3.6 Group Analysis of Post-translational Modifications in the Rapeseed Oil Group

The post-translational modifications of the rapeseed oil group were compared with those of the control group. The differential modification screening criteria were FC ≥ 1.5 or ≤ 0.67, and a two-tailed paired t-test with P < 0.05. The results showed that, compared to the control group, a total of 50 differential modifications were identified in the rapeseed oil group, with 38 down-regulated and 12 up-regulated. The differentially modified proteins were arranged in ascending order of FC, and the detailed information is listed in Appendix Table 10.

Among the proteins with differential modifications showing an up- or down-regulation of more than 10-fold: P07632, namely Superoxide dismutase (FC = 0, P = 2.95E-02), can neutralize superoxide free radicals and protect organisms from oxidative stress, playing an important role in the antioxidant process^[66]^. P47853, namely Biglycan (FC = 0.045, P = 4.47E-02), can reduce high-fat-diet-induced obesity and improve glucose tolerance in mice^[67]^.

## 4 Conclusion

After rats consumed edible oil for one week, significant changes occurred in the urinary proteome. Moreover, different types of edible oils had remarkable differences in their effects on the urinary proteome of rats, demonstrating a comprehensive impact on the rat body. Compared with the urinary proteome itself, the changes in post-translational modifications of the proteome were relatively small, indicating that the intake of edible oils in the short term had a relatively limited impact on protein post-translational modifications.

## References

[1] Simopoulos AP. The importance of the omega-6/omega-3 fatty acid ratio in cardiovascular disease and other chronic diseases. Exp Biol Med (Maywood). 2008 Jun;233(6):674–88. doi: 10.3181/0711-MR-311. Epub 2008 Apr 11. PMID: 18408140.

[2] Kris-Etherton PM. AHA science advisory: monounsaturated fatty acids and risk of cardiovascular disease. J Nutr. 1999 Dec;129(12):2280–4. doi: 10.1093/jn/129.12.2280. PMID: 10573564.

[3] Xia M, Zhong Y, Peng Y, Qian C. Olive oil consumption and risk of cardiovascular disease and all-cause mortality: A meta-analysis of prospective cohort studies. Front Nutr. 2022 Oct 18;9:1041203. doi: 10.3389/fnut.2022.1041203. PMID: 36330142; PMCID: PMC9623257.

[4] Hu FB, Stampfer MJ, Manson JE, Ascherio A, Colditz GA, Speizer FE, Hennekens CH, Willett WC. Dietary saturated fats and their food sources in relation to the risk of coronary heart disease in women. Am J Clin Nutr. 1999 Dec;70(6):1001–8. doi: 10.1093/ajcn/70.6.1001. PMID: 10584044.

[5] Yan S, Liu S, Qu J, Li X, Hu J, Zhang L, Liu X, Li X, Wang X, Wen L, Wang J. A Lard and Soybean Oil Mixture Alleviates Low-Fat-High-Carbohydrate Diet-Induced Nonalcoholic Fatty Liver Disease in Mice. Nutrients. 2022 Jan 27;14(3):560. doi: 10.3390/nu14030560. PMID: 35276916; PMCID: PMC8840387.

[6] Li H, Zhu Y, Zhao F, Song S, Li Y, Xu X, Zhou G, Li C. Fish oil, lard and soybean oil differentially shape gut microbiota of middle-aged rats. Sci Rep. 2017 Apr 11;7(1):826. doi: 10.1038/s41598-017-00969-0. Erratum in: Sci Rep. 2017 Aug 10;7(1):7738. doi: 10.1038/s41598-017-04862-8. PMID: 28400577; PMCID: PMC5429820.

[7] Mozaffarian D, Katan MB, Ascherio A, Stampfer MJ, Willett WC. Trans fatty acids and cardiovascular disease. N Engl J Med. 2006 Apr 13;354(15):1601–13. doi: 10.1056/NEJMra054035. PMID: 16611951.

[8] Fallah Z, Vasmehjani AA, Aghaei S, Amiri M, Raeisi-Dekordi H, Moghtaderi F, Zimorovat A, Yazd EF, Madadizadeh F, Khayyatzadeh SS, Salehi-Abargouei A. Cardiometabolic risk factors are affected by interaction between FADS1 rs174556 variant and dietary vegetable oils in patients with diabetes: a randomized controlled trial. Sci Rep. 2024 Nov 11;14(1):27531. doi: 10.1038/s41598-024-78294-6. PMID: 39528535; PMCID: PMC11555249.

[9] Rudman N, Gornik O, Lauc G. Altered N-glycosylation profiles as potential biomarkers and drug targets in diabetes. FEBS Lett. 2019 Jul;593(13):1598–1615. doi: 10.1002/1873-3468.13495. Epub 2019 Jun 28. PMID: 31215021.

[10] Sen, Anuhna et al. “Scaffolding Protein IQ Motif Containing GTPase Activating Protein 2 Regulates Liver Metabolic Homeostasis.” The FASEB journal. 34.S1 (2020): 1–1. Web.

[11] Shetty S, Copeland PR. Molecular mechanism of selenoprotein P synthesis. Biochim Biophys Acta Gen Subj. 2018 Nov;1862(11):2506–2510. doi: 10.1016/j.bbagen.2018.04.011. Epub 2018 Apr 12. PMID: 29656121; PMCID: PMC6188828.

[12] Christians ES, Ishiwata T, Benjamin IJ. Small heat shock proteins in redox metabolism: implications for cardiovascular diseases. Int J Biochem Cell Biol. 2012 Oct;44(10):1632–45. doi: 10.1016/j.biocel.2012.06.006. Epub 2012 Jun 15. PMID: 22710345; PMCID: PMC3412898.

[13] Grewal R, Reutzel M, Dilberger B, Hein H, Zotzel J, Marx S, Tretzel J, Sarafeddinov A, Fuchs C, Eckert GP. Purified oleocanthal and ligstroside protect against mitochondrial dysfunction in models of early Alzheimer’s disease and brain ageing. Exp Neurol. 2020 Jun;328:113248. doi: 10.1016/j.expneurol.2020.113248. Epub 2020 Feb 19. PMID: 32084452.

[14] Scholz A, Plate KH, Reiss Y. Angiopoietin-2: a multifaceted cytokine that functions in both angiogenesis and inflammation. Ann N Y Acad Sci. 2015 Jul;1347:45–51. doi: 10.1111/nyas.12726. Epub 2015 Mar 13. PMID: 25773744.

[15] Jiang P, Ning J, Yu W, Rao T, Ruan Y, Cheng F. FLRT2 suppresses bladder cancer progression through inducing ferroptosis. J Cell Mol Med. 2024 Mar;28(5):e17855. doi: 10.1111/jcmm.17855. Epub 2023 Jul 21. PMID: 37480224; PMCID: PMC10902570.

[16] Mishima M, Sakai Y, Itoh N, Kamiya H, Furuichi M, Takahashi M, Yamagata Y, Iwai S, Nakabeppu Y, Shirakawa M. Structure of human MTH1, a Nudix family hydrolase that selectively degrades oxidized purine nucleoside triphosphates. J Biol Chem. 2004 Aug 6;279(32):33806–15. doi: 10.1074/jbc.M402393200. Epub 2004 May 7. PMID: 15133035.

[17] Barbalace MC, Zallocco L, Beghelli D, Ronci M, Scortichini S, Digiacomo M, Macchia M, Mazzoni MR, Fiorini D, Lucacchini A, Hrelia S, Giusti L, Angeloni C. Antioxidant and Neuroprotective Activity of Extra Virgin Olive Oil Extracts Obtained from Quercetano Cultivar Trees Grown in Different Areas of the Tuscany Region (Italy). Antioxidants (Basel). 2021 Mar 10;10(3):421. doi: 10.3390/antiox10030421. PMID: 33801925; PMCID: PMC8000409.

[18] Villareal MO, Sasaki K, Margout D, Savry C, Almaksour Z, Larroque M, Isoda H. Neuroprotective effect of Picholine virgin olive oil and its hydroxycinnamic acids component against β-amyloid-induced toxicity in SH-SY5Y neurotypic cells. Cytotechnology. 2016 Dec;68(6):2567–2578. doi: 10.1007/s10616-016-9980-3. Epub 2016 May 7. PMID: 27155966; PMCID: PMC5101328.

[19] Lambert de Malezieu M, Courtel P, Sleno L, Abasq ML, Ramassamy C. Synergistic properties of bioavailable phenolic compounds from olive oil: electron transfer and neuroprotective properties. Nutr Neurosci. 2021 Sep;24(9):660–673. doi: 10.1080/1028415X.2019.1666480. Epub 2019 Oct 9. PMID: 31595838.

[20] Catalán Ú, Rubió L, López de Las Hazas MC, Herrero P, Nadal P, Canela N, Pedret A, Motilva MJ, Solà R. Hydroxytyrosol and its complex forms (secoiridoids) modulate aorta and heart proteome in healthy rats: Potential cardio-protective effects. Mol Nutr Food Res. 2016 Oct;60(10):2114–2129. doi: 10.1002/mnfr.201600052. Epub 2016 Jun 1. PMID: 27125338.

[21] Gholamalizadeh H, Ensan B, Karav S, Jamialahmadi T, Sahebkar A. Regulatory effects of statins on CCL2/CCR2 axis in cardiovascular diseases: new insight into pleiotropic effects of statins. J Inflamm (Lond). 2024 Dec 18;21(1):51. doi: 10.1186/s12950-024-00420-y. PMID: 39696507; PMCID: PMC11658147.

[22] León A, Aparicio GI, Scorticati C. Neuronal Glycoprotein M6a: An Emerging Molecule in Chemical Synapse Formation and Dysfunction. Front Synaptic Neurosci. 2021 May 4;13:661681. doi: 10.3389/fnsyn.2021.661681. PMID: 34017241; PMCID: PMC8129562.

[23] Solovyev N. Selenoprotein P and its potential role in Alzheimer’s disease. Hormones (Athens). 2020 Mar;19(1):73–79. doi: 10.1007/s42000-019-00112-w. Epub 2019 Jun 27. PMID: 31250406.

[24] Menghini R, Casagrande V, Menini S, Marino A, Marzano V, Hribal ML, Gentileschi P, Lauro D, Schillaci O, Pugliese G, Sbraccia P, Urbani A, Lauro R, Federici M. TIMP3 overexpression in macrophages protects from insulin resistance, adipose inflammation, and nonalcoholic fatty liver disease in mice. Diabetes. 2012 Feb;61(2):454–62. doi: 10.2337/db11-0613. Epub 2012 Jan 6. PMID: 22228717; PMCID: PMC3266402.

[25] Herrera-Marcos LV, Lou-Bonafonte JM, Martinez-Gracia MV, Arnal C, Navarro MA, Osada J. Prenylcysteine oxidase 1, a pro-oxidant enzyme of low density lipoproteins. Front Biosci (Landmark Ed). 2018 Jan 1;23(6):1020–1037. doi: 10.2741/4631. PMID: 28930587.

[26] Tian X, Yan C, Han Y. Cellular Repressor of E1A-stimulated Genes, A New Potential Therapeutic Target for Atherosclerosis. Curr Drug Targets. 2017;18(15):1800–1804. doi: 10.2174/1389450117666161026111250. PMID: 27784214.

[27] Liu Y, Zhao W, Gu G, Lu L, Feng J, Guo Q, Ding Z. Palmitoyl-protein thioesterase 1 (PPT1): an obesity-induced rat testicular marker of reduced fertility. Mol Reprod Dev. 2014 Jan;81(1):55–65. doi: 10.1002/mrd.22281. Epub 2013 Dec 3. PMID: 24302477.

[28] Jin DZ, Mao LM, Wang JQ. The Role of Extracellular Signal-Regulated Kinases (ERK) in the Regulation of mGlu5 Receptors in Neurons. J Mol Neurosci. 2018 Dec;66(4):629–638. doi: 10.1007/s12031-018-1193-0. Epub 2018 Nov 15. PMID: 30430306; PMCID: PMC6312115.

[29] Wright NT, Cannon BR, Zimmer DB, Weber DJ. S100A1: Structure, Function, and Therapeutic Potential. Curr Chem Biol. 2009 May 1;3(2):138–145. doi: 10.2174/187231309788166460. PMID: 19890475; PMCID: PMC2771873.

[30] Zhang L, Sun C, Jin Y, Gao K, Shi X, Qiu W, Ma C, Zhang L. Dickkopf 3 (Dkk3) Improves Amyloid-β Pathology, Cognitive Dysfunction, and Cerebral Glucose Metabolism in a Transgenic Mouse Model of Alzheimer’s Disease. J Alzheimers Dis. 2017;60(2):733–746. doi: 10.3233/JAD-161254. PMID: 28922151.

[31] Lizarraga F, Espinosa M, Ceballos-Cancino G, Vazquez-Santillan K, Bahena-Ocampo I, Schwarz-Cruz Y Celis A, Vega-Gordillo M, Garcia Lopez P, Maldonado V, Melendez-Zajgla J. Tissue inhibitor of metalloproteinases-4 (TIMP-4) regulates stemness in cervical cancer cells. Mol Carcinog. 2016 Dec;55(12):1952–1961. doi: 10.1002/mc.22442. Epub 2015 Nov 30. PMID: 26618609.

[32] Guo Z, Zhao Y, Wu Y, Zhang Y, Wang R, Liu W, Zhang C, Yang X. Cellular retinol-binding protein 1: a therapeutic and diagnostic tumor marker. Mol Biol Rep. 2023 Feb;50(2):1885–1894. doi: 10.1007/s11033-022-08179-2. Epub 2022 Dec 14. PMID: 36515825.

[33] Zhong M, Kawaguchi R, Ter-Stepanian M, Kassai M, Sun H. Vitamin A transport and the transmembrane pore in the cell-surface receptor for plasma retinol binding protein. PLoS One. 2013 Nov 1;8(11):e73838. doi: 10.1371/journal.pone.0073838. PMID: 24223695; PMCID: PMC3815300.

[34] Yang H, Hu B, Wang X, Chen W, Zhou H. The effects of hyaluronan and proteoglycan link protein 1 (HAPLN1) in ameliorating spinal cord injury mediated by Nrf2. Biotechnol Appl Biochem. 2024 Aug;71(4):929–939. doi: 10.1002/bab.2587. Epub 2024 Apr 12. PMID: 38607990.

[35] Brown CW, Chhoy P, Mukhopadhyay D, Karner ER, Mercurio AM. Targeting prominin2 transcription to overcome ferroptosis resistance in cancer. EMBO Mol Med. 2021 Aug 9;13(8):e13792. doi: 10.15252/emmm.202013792. Epub 2021 Jul 5. PMID: 34223704; PMCID: PMC8350900.

[36] Luo J, Xie M, Peng C, Ma Y, Wang K, Lin G, Yang H, Chen T, Liu Q, Zhang G, Lin H, Ji Z. Protein disulfide isomerase A6 promotes the repair of injured nerve through interactions with spastin. Front Mol Neurosci. 2022 Aug 24;15:950586. doi: 10.3389/fnmol.2022.950586. PMID: 36090256; PMCID: PMC9449696.

[37] Pucci L, Perozzi S, Cimadamore F, Orsomando G, Raffaelli N. Tissue expression and biochemical characterization of human 2-amino 3-carboxymuconate 6-semialdehyde decarboxylase, a key enzyme in tryptophan catabolism. FEBS J. 2007 Feb;274(3):827–40. doi: 10.1111/j.1742-4658.2007.05635.x. PMID: 17288562.

[38] Adamson SE, Griffiths R, Moravec R, Senthivinayagam S, Montgomery G, Chen W, Han J, Sharma PR, Mullins GR, Gorski SA, Cooper JA, Kadl A, Enfield K, Braciale TJ, Harris TE, Leitinger N. Disabled homolog 2 controls macrophage phenotypic polarization and adipose tissue inflammation. J Clin Invest. 2016 Apr 1;126(4):1311–22. doi: 10.1172/JCI79590. Epub 2016 Feb 29. PMID: 26927671; PMCID: PMC4811113.

[39] Ermolova N, Kramerova I, Spencer MJ. Autolytic activation of calpain 3 proteinase is facilitated by calmodulin protein. J Biol Chem. 2015 Jan 9;290(2):996–1004. doi: 10.1074/jbc.M114.588780. Epub 2014 Nov 11. PMID: 25389288; PMCID: PMC4294526.

[40] Williams AS, Kang L, Zheng J, Grueter C, Bracy DP, James FD, Pozzi A, Wasserman DH. Integrin α1-null mice exhibit improved fatty liver when fed a high fat diet despite severe hepatic insulin resistance. J Biol Chem. 2015 Mar 6;290(10):6546–57. doi: 10.1074/jbc.M114.615716. Epub 2015 Jan 15. PMID: 25593319; PMCID: PMC4358288.

[41] Bourebaba L, Marycz K. Pathophysiological Implication of Fetuin-A Glycoprotein in the Development of Metabolic Disorders: A Concise Review. J Clin Med. 2019 Nov 21;8(12):2033. doi: 10.3390/jcm8122033. PMID: 31766373; PMCID: PMC6947209.

[42] Xu J, He M, Wang W, Hou J, Chen X, Ding X, Zhang J. siRNA-mediated Eppin testicular silencing causes changes in sperm motility and calcium currents in mice. Reprod Biol. 2021 Jun;21(2):100485. doi: 10.1016/j.repbio.2021.100485. Epub 2021 Feb 16. PMID: 33607572.

[43] Dauvilliers Y, Tafti M, Landolt HP. Catechol-O-methyltransferase, dopamine, and sleep-wake regulation. Sleep Med Rev. 2015 Aug;22:47–53. doi: 10.1016/j.smrv.2014.10.006. Epub 2014 Oct 27. PMID: 25466290.

[44] Brody MJ, Vanhoutte D, Schips TG, Boyer JG, Bakshi CV, Sargent MA, York AJ, Molkentin JD. Defective Flux of Thrombospondin-4 through the Secretory Pathway Impairs Cardiomyocyte Membrane Stability and Causes Cardiomyopathy. Mol Cell Biol. 2018 Jun 28;38(14):e00114–18. doi: 10.1128/MCB.00114-18. PMID: 29712757; PMCID: PMC6024163.

[45] Geginat J. Introduction to the Special Issue: Interleukin-10 “The surprising twists and turns of an anti-inflammatory cytokine on its way to the clinic”. Semin Immunol. 2019 Aug;44:101343. doi: 10.1016/j.smim.2019.101343. Epub 2019 Nov 6. PMID: 31706854.

[46] He JQ, Shumansky K, Zhang X, Connett JE, Anthonisen NR, Sandford AJ. Polymorphisms of interleukin-10 and its receptor and lung function in COPD. Eur Respir J. 2007 Jun;29(6):1120–6. doi: 10.1183/09031936.00002907. Epub 2007 Mar 1. PMID: 17331973.

[47] Zhao C, Rispe C, Nabity PD. Secretory RING finger proteins function as effectors in a grapevine galling insect. BMC Genomics. 2019 Dec 3;20(1):923. doi: 10.1186/s12864-019-6313-x. PMID: 31795978; PMCID: PMC6892190.

[48] Wang K, Kim MK, Di Caro G, Wong J, Shalapour S, Wan J, Zhang W, Zhong Z, Sanchez-Lopez E, Wu LW, Taniguchi K, Feng Y, Fearon E, Grivennikov SI, Karin M. Interleukin-17 receptor a signaling in transformed enterocytes promotes early colorectal tumorigenesis. Immunity. 2014 Dec 18;41(6):1052–63. doi: 10.1016/j.immuni.2014.11.009. Epub 2014 Nov 25. PMID: 25526314; PMCID: PMC4272447.

[49] O’Hehir ZD, Lynch T, O’Neill S, March L, Xue M. Endothelial Protein C Receptor and Its Impact on Rheumatic Disease. J Clin Med. 2024 Mar 31;13(7):2030. doi: 10.3390/jcm13072030. PMID: 38610795; PMCID: PMC11012567.

[50] Walker DG, Lue LF. Understanding the neurobiology of CD200 and the CD200 receptor: a therapeutic target for controlling inflammation in human brains? Future Neurol. 2013 May;8(3):10.2217/fnl.13.14. doi: 10.2217/fnl.13.14. PMID: 24198718; PMCID: PMC3815586.

[51] Feng Y, Jiang H, Li G, He G, Li X. Decreased expression of protein *O*-linked mannose β-1,2-*N*-acetylglucosaminyltransferase 1 contributes to Alzheimer’s disease-like pathologies. J Neurophysiol. 2022 Apr 1;127(4):1067–1074. doi: 10.1152/jn.00362.2021. Epub 2022 Mar 23. PMID: 35320023.

[52] Liu Y, Jiang S, Li Q, Kong Y. [Advances of Kunitz-type serine protease inhibitors]. Sheng Wu Gong Cheng Xue Bao. 2021 Nov 25;37(11):3988–4000. Chinese. doi: 10.13345/j.cjb.200802. PMID: 34841799.

[53] Du H, Zhou Y, Suo Y, Liang X, Chai B, Duan R, Huang X, Li Q. CCN1 accelerates re-epithelialization by promoting keratinocyte migration and proliferation during cutaneous wound healing. Biochem Biophys Res Commun. 2018 Nov 10;505(4):966–972. doi: 10.1016/j.bbrc.2018.09.001. Epub 2018 Oct 22. PMID: 30361094.

[54] Hillyer LM, Woodward B. A comparison of the capacity of six cold-pressed plant oils to support development of acquired immune competence in the weanling mouse: superiority of low-linoleic-acid oils. Br J Nutr. 2002 Aug;88(2):171–81. doi: 10.1079/BJNBJN2002602. PMID: 12144720.

[55] Tiwari S, Atluri V, Kaushik A, Yndart A, Nair M. Alzheimer’s disease: pathogenesis, diagnostics, and therapeutics. Int J Nanomedicine. 2019 Jul 19;14:5541–5554. doi: 10.2147/IJN.S200490. PMID: 31410002; PMCID: PMC6650620.

[56] Hashempour-Baltork F, Farshi P, Mirza Alizadeh A, Eskandarzadeh S, Abedinzadeh S, Azadmard-Damirchi S, Torbati M. Effect of Refined Edible Oils on Neurodegenerative Disorders. Adv Pharm Bull. 2023 Jul;13(3):461–468. doi: 10.34172/apb.2023.060. Epub 2022 Nov 2. PMID: 37646051; PMCID: PMC10460797.

[57] Urbányi Z, Forrai E, Sárvári M, Likó I, Illés J, Pázmány T. Glycosaminoglycans inhibit neurodegenerative effects of serum amyloid P component in vitro. Neurochem Int. 2005 May;46(6):471–7. doi: 10.1016/j.neuint.2004.12.001. PMID: 15769549.

[58] Wang J, Yu JT, Tan L. PLD3 in Alzheimer’s disease. Mol Neurobiol. 2015 Apr;51(2):480–6. doi: 10.1007/s12035-014-8779-5. Epub 2014 Jun 17. PMID: 24935720.

[59] Kobayashi R, Nakagomi Y, Shimura Y, Mochizuki M, Kobayashi K, Sugita K, Ohyama K. Expression of stanniocalcin-1 in gastrointestinal tracts of neonatal and mature rats. Biochem Biophys Res Commun. 2009 Nov 20;389(3):478–83. doi: 10.1016/j.bbrc.2009.08.169. Epub 2009 Sep 2. PMID: 19732741.

[60] Peters T Jr. Serum albumin. Adv Clin Chem. 1970;13:37–111. doi: 10.1016/s0065-2423(08)60385-6. PMID: 4919632.

[61] Procter M, Wolf B, Crockett DK, Mao R. The Biotinidase Gene Variants Registry: A Paradigm Public Database. G3 (Bethesda). 2013 Apr 9;3(4):727–731. doi: 10.1534/g3.113.005835. PMID: 23550138; PMCID: PMC3618359.

[62] Urbányi Z, Forrai E, Sárvári M, Likó I, Illés J, Pázmány T. Glycosaminoglycans inhibit neurodegenerative effects of serum amyloid P component in vitro. Neurochem Int. 2005 May;46(6):471–7. doi: 10.1016/j.neuint.2004.12.001. PMID: 15769549.

[63] Althubiti M, Elzubier M, Alotaibi GS, Althubaiti MA, Alsadi HH, Alhazmi ZA, Alghamdi F, El-Readi MZ, Almaimani R, Babakr A. Beta 2 microglobulin correlates with oxidative stress in elderly. Exp Gerontol. 2021 Jul 15;150:111359. doi: 10.1016/j.exger.2021.111359. Epub 2021 Apr 24. PMID: 33905876.

[64] Jaganjac M, Milkovic L, Sunjic SB, Zarkovic N. The NRF2, Thioredoxin, and Glutathione System in Tumorigenesis and Anticancer Therapies. Antioxidants (Basel). 2020 Nov 19;9(11):1151. doi: 10.3390/antiox9111151. PMID: 33228209; PMCID: PMC7699519.

[65] Wolf MTF, Zhang J, Nie M. Uromodulin in mineral metabolism. Curr Opin Nephrol Hypertens. 2019 Sep;28(5):481–489. doi: 10.1097/MNH.0000000000000522. PMID: 31205055; PMCID: PMC6764599.

[66] Canada AT, Calabrese EJ. Superoxide dismutase: its role in xenobiotic detoxification. Pharmacol Ther. 1989;44(2):285–95. doi: 10.1016/0163-7258(89)90068-5. PMID: 2519345.

[67] Chung I, Kim SA, Kim S, Lee JO, Park CY, Lee J, Kang J, Lee JY, Seo I, Lee HJ, Han JA, Kang MJ, Lim E, Kim SJ, Wu SW, Oh JY, Chung JH, Kim EK, Kim HS, Shin MJ. Biglycan reduces body weight by regulating food intake in mice and improves glucose metabolism through AMPK/AKT dual pathways in skeletal muscle. FASEB J. 2021 Aug;35(8):e21794. doi: 10.1096/fj.202002039RR. PMID: 34314059.

