## Appendix for "Effects of One-week Intake of Different Edible Oils on the Urinary Proteome of Rats"

Appendix Table 1 Differentially Expressed Proteins in the Urinary Proteome of the Olive Oil Group and the Control Group Screened with FC≥1.5 or ≤0.67, P < 0.05

| Uniprot ID | Protein name | Fold change | P-value | Trend |
| --- | --- | --- | --- | --- |
| F1LMP1 | Alpha-1,3-mannosyl-glycoprotein 4-beta-N-acetylglucosaminyltransferase A | 0.000 | 3.81E-02 | ↓ |
| Q5XIF6 | Tubulin alpha-4A chain | 0.020 | 3.84E-02 | ↓ |
| P18614 | Integrin alpha-1 | 0.047 | 2.81E-02 | ↓ |
| F1LW74 | IQ motif containing GTPase activating protein 2 | 0.094 | 2.22E-02 | ↓ |
| A0A0G2K0V8 | Nuclear distribution C, dynein complex regulator | 0.104 | 2.73E-02 | ↓ |
| M0R7T1 | Phosphatase and actin regulator 4 | 0.180 | 2.94E-02 | ↓ |
| A0A0G2JSZ2 | Seminal vesicle secretory protein 4 | 0.195 | 4.87E-02 | ↓ |
| P25236 | Selenoprotein P | 0.213 | 1.71E-03 | ↓ |
| F1LQM1 | Alpha-2u-globulin | 0.267 | 4.55E-02 | ↓ |
| Q8R2H2 | Integrin beta-3 | 0.269 | 4.35E-03 | ↓ |
| P34058 | Heat shock protein HSP 90-beta | 0.318 | 8.62E-03 | ↓ |
| B1WBS5 | Sodium/mannose cotransporter SLC5A10 | 0.349 | 4.09E-02 | ↓ |
| Q9JJI3 | Major urinary protein 4 | 0.386 | 3.35E-02 | ↓ |
| P16573 | Carcinoembryonic antigen-related cell adhesion molecule 1 | 0.398 | 2.46E-02 | ↓ |
| F8WG88 | Follistatin-related protein 1 | 0.404 | 4.87E-02 | ↓ |
| A0A0G2JXZ9 | protein-tyrosine-phosphatase | 0.412 | 7.16E-03 | ↓ |
| G3V9J1 | Murinoglobulin 2 | 0.467 | 4.89E-02 | ↓ |
| A0A0G2JTX6 | C-type lectin domain containing 5A | 1.574 | 3.76E-02 | ↑ |
| M0R7Q2 | Ig-like domain-containing protein | 1.645 | 4.40E-02 | ↑ |
| Q5U2Q3 | Ester hydrolase C11orf54 homolog | 1.835 | 1.96E-02 | ↑ |
| Q9ESS6 | Basal cell adhesion molecule | 1.840 | 3.93E-02 | ↑ |
| F1M663 | Ig-like domain-containing protein | 2.041 | 3.09E-03 | ↑ |
| P70490 | Lactadherin | 2.055 | 4.93E-02 | ↑ |
| P04785 | Protein disulfide-isomerase | 2.093 | 3.12E-02 | ↑ |
| A0A0G2JXT8 | Filamin B | 2.100 | 4.10E-02 | ↑ |
| P07150 | Annexin A1 | 2.313 | 3.33E-02 | ↑ |
| D3ZQP6 | Semaphorin 7A | 2.331 | 2.94E-02 | ↑ |
| F7F3I7 | Semaphorin 4A | 2.381 | 4.06E-02 | ↑ |
| Q5U2V4 | Phospholipase B-like 1 | 2.407 | 2.97E-02 | ↑ |
| A0A0G2JZN1 | deleted | 2.460 | 2.72E-02 | ↑ |
| Q5BK81 | Prostaglandin reductase 2 | 2.481 | 6.86E-03 | ↑ |
| F1LSV0 | Semaphorin 4B | 2.523 | 1.37E-02 | ↑ |
| F1LPC6 | Carboxypeptidase D | 2.532 | 9.34E-03 | ↑ |
| D3ZHA1 | Beta-1,4-glucuronyltransferase 1 | 2.608 | 4.44E-02 | ↑ |
| D4ABU7 | Triggering receptor expressed on myeloid cells 1 | 2.614 | 4.02E-02 | ↑ |
| A0A140TAI2 | Biotinidase | 2.642 | 1.83E-02 | ↑ |
| A0A0G2JY48 | receptor protein-tyrosine kinase | 2.778 | 3.02E-04 | ↑ |
| M0RDL2 | deleted | 2.869 | 2.35E-02 | ↑ |
| P09034 | Argininosuccinate synthase | 2.873 | 4.08E-02 | ↑ |
| G3V647 | Pyridoxal kinase | 2.958 | 3.32E-02 | ↑ |
| P15429 | Beta-enolase | 2.983 | 3.95E-02 | ↑ |
| B1WBN7 | FAM20C | 2.996 | 3.27E-02 | ↑ |
| Q6AYE5 | Out at first protein homolog | 3.081 | 1.77E-02 | ↑ |
| Q00657 | Chondroitin sulfate proteoglycan 4 | 3.180 | 1.19E-02 | ↑ |
| Q66HG4 | Galactose mutarotase | 3.231 | 2.82E-02 | ↑ |
| P80254 | D-dopachrome decarboxylase | 3.465 | 4.00E-02 | ↑ |
| D3ZBS2 | Inter-alpha-trypsin inhibitor heavy chain H3 | 3.466 | 4.06E-02 | ↑ |
| B5DEI2 | Amine oxidase | 3.502 | 3.78E-02 | ↑ |
| Q6P7S1 | Acid ceramidase | 3.578 | 1.69E-02 | ↑ |
| Q80ZA3 | Serine | 3.626 | 1.73E-02 | ↑ |
| M0RAM5 | Glutathione peroxidase | 3.773 | 2.25E-03 | ↑ |
| Q66H12 | Alpha-N-acetylgalactosaminidase | 3.783 | 1.41E-02 | ↑ |
| P04905 | Glutathione S-transferase Mu 1 | 3.885 | 9.36E-03 | ↑ |
| Q64361 | Latexin | 3.898 | 2.47E-02 | ↑ |
| Q6JE36 | Protein NDRG1 | 4.046 | 1.28E-02 | ↑ |
| F1MAN8 | Laminin subunit alpha 5 | 4.061 | 4.57E-02 | ↑ |
| Q80WD1 | Reticulon-4 receptor-like 2 | 4.067 | 2.21E-02 | ↑ |
| B0BN46 | Grhpr protein | 4.087 | 3.79E-02 | ↑ |
| Q9R1T3 | Cathepsin Z | 4.163 | 1.10E-02 | ↑ |
| M0R3V4 | Myeloid-derived growth factor | 4.195 | 5.85E-03 | ↑ |
| P19804 | Nucleoside diphosphate kinase B | 4.259 | 4.81E-02 | ↑ |
| Q64573 | Liver carboxylesterase 1F | 4.302 | 4.58E-02 | ↑ |
| B1H232 | Immunoglobulin superfamily containing leucine-rich repeat | 4.366 | 4.88E-03 | ↑ |
| A0A0G2JZS9 | Ig-like domain-containing protein | 4.371 | 4.73E-02 | ↑ |
| P47853 | Biglycan | 4.378 | 4.97E-02 | ↑ |
| A2IBE0 | Membrane-bound carbonic anhydrase 14 | 4.589 | 1.39E-03 | ↑ |
| A0A0G2JXN4 | Glutathione peroxidase | 4.644 | 3.44E-02 | ↑ |
| A0A0G2JV12 | Collagen type XV alpha 1 chain | 4.660 | 4.47E-02 | ↑ |
| Q4FZV0 | Beta-mannosidase | 4.713 | 9.92E-04 | ↑ |
| Q32KJ6 | N-acetylgalactosamine-6-sulfatase | 4.750 | 1.15E-02 | ↑ |
| P22734 | Catechol O-methyltransferase | 4.764 | 9.59E-04 | ↑ |
| O35760 | Isopentenyl-diphosphate Delta-isomerase 1 | 4.936 | 4.64E-02 | ↑ |
| Q675A5 | Lysosomal phospholipase A and acyltransferase | 5.090 | 2.79E-02 | ↑ |
| Q99M75 | Reticulon-4 receptor | 5.202 | 1.61E-02 | ↑ |
| Q920A6 | Retinoid-inducible serine carboxypeptidase | 5.230 | 1.80E-02 | ↑ |
| F1LP42 | Hedgehog protein | 5.268 | 4.94E-03 | ↑ |
| Q5PPH0 | Enolase-phosphatase E1 | 5.578 | 2.77E-02 | ↑ |
| Q641Z7 | Cyclic GMP-AMP phosphodiesterase SMPDL3A | 5.628 | 4.19E-02 | ↑ |
| A0A0H2UHH2 | Amyloid P component, serum | 5.642 | 1.99E-02 | ↑ |
| P45479 | Palmitoyl-protein thioesterase 1 | 5.755 | 1.20E-02 | ↑ |
| D4ACR1 | RCG64219-like | 6.063 | 4.08E-02 | ↑ |
| Q9EQV6 | Tripeptidyl-peptidase 1 | 6.607 | 2.50E-02 | ↑ |
| F1M7B3 | Ig-like domain-containing protein | 6.880 | 1.68E-02 | ↑ |
| Q562C9 | Acireductone dioxygenase | 7.269 | 2.55E-02 | ↑ |
| G3V722 | Beta-1,4-galactosyltransferase | 7.724 | 4.13E-02 | ↑ |
| G3V862 | Angiopoietin-like 2 | 7.891 | 1.18E-02 | ↑ |
| G3V7N8 | Galectin 9 | 7.948 | 4.30E-03 | ↑ |
| Q5I0D5 | Phospholysine phosphohistidine inorganic pyrophosphate phosphatase | 8.140 | 3.59E-02 | ↑ |
| D3ZTV3 | Leucine-rich repeat transmembrane protein FLRT2 | 8.891 | 5.15E-03 | ↑ |
| F1M091 | Kallikrein m | 9.236 | 1.33E-02 | ↑ |
| D3ZP44 | SLIT and NTRK-like family, member 6 | 9.527 | 1.06E-02 | ↑ |
| Q5FVH2 | 5'-3' exonuclease PLD3 | 9.588 | 7.19E-03 | ↑ |
| P53369 | Oxidized purine nucleoside triphosphate hydrolase | 10.262 | 2.83E-03 | ↑ |
| F1M0B7 | deleted | 11.606 | 4.74E-02 | ↑ |

Appendix Table 2 Differentially Expressed Proteins in the Urinary Proteome of the Butter Group and the Control Group Screened with FC≥1.5 or ≤0.67, P < 0.05

| Uniprot ID | Protein name | Fold change | P-value | Trend |
| --- | --- | --- | --- | --- |
| P14844 | C-C motif chemokine 2 | 0.190 | 2.18E-02 | ↓ |
| M0R7T1 | Phosphatase and actin regulator 4 | 0.239 | 4.91E-02 | ↓ |
| A0A0G2JXY4 | Carbonic anhydrase 15 | 0.258 | 4.02E-02 | ↓ |
| Q812E9 | Neuronal membrane glycoprotein M6-a | 0.277 | 2.56E-02 | ↓ |
| P25236 | Selenoprotein P | 0.340 | 2.72E-04 | ↓ |
| P34058 | Heat shock protein HSP 90-beta | 0.351 | 1.10E-02 | ↓ |
| Q9JJ40 | Na(+)/H(+) exchange regulatory cofactor NHE-RF3 | 0.364 | 1.70E-02 | ↓ |
| B4F7A5 | Cd99 protein | 0.412 | 1.17E-02 | ↓ |
| A0A0G2JXZ9 | protein-tyrosine-phosphatase | 0.415 | 1.41E-02 | ↓ |
| F8WG88 | Follistatin-related protein 1 | 0.484 | 3.68E-02 | ↓ |
| Q5M876 | N-acyl-aromatic-L-amino acid amidohydrolase | 0.508 | 1.56E-02 | ↓ |
| Q8R2H2 | Integrin beta-3 | 0.509 | 1.98E-02 | ↓ |
| D3Z9U8 | S100 calcium binding protein A7 like 2 | 0.542 | 2.15E-02 | ↓ |
| G3V6A0 | Platelet-derived growth factor receptor alpha | 0.603 | 3.98E-02 | ↓ |
| Q5HZW5 | CD320 antigen | 0.668 | 2.51E-02 | ↓ |
| P04785 | Protein disulfide-isomerase | 1.600 | 1.80E-02 | ↑ |
| F1LPC6 | Carboxypeptidase D | 1.673 | 2.01E-02 | ↑ |
| D3ZZ44 | Extracellular leucine-rich repeat and fibronectin type III domain containing 1 | 1.713 | 1.37E-03 | ↑ |
| D3ZQP6 | Semaphorin 7A | 1.740 | 1.47E-02 | ↑ |
| Q00657 | Chondroitin sulfate proteoglycan 4 | 1.769 | 4.27E-02 | ↑ |
| A0A0G2JTX5 | Dipeptidyl peptidase 4 | 1.817 | 3.56E-02 | ↑ |
| M0RDA4 | receptor protein-tyrosine kinase | 1.865 | 2.60E-02 | ↑ |
| P47820 | Angiotensin-converting enzyme | 1.969 | 2.16E-02 | ↑ |
| Q5M843 | 2-oxoglutarate and iron-dependent oxygenase domain-containing protein 3 | 2.027 | 3.65E-02 | ↑ |
| G3V989 | EPH-related receptor tyrosine kinase ligand 7 | 2.033 | 1.78E-02 | ↑ |
| Q6JE36 | Protein NDRG1 | 2.076 | 3.64E-02 | ↑ |
| G3V8P3 | Cadherin, EGF LAG seven-pass G-type receptor 2 | 2.091 | 2.43E-02 | ↑ |
| M0R3V4 | Myeloid-derived growth factor | 2.139 | 1.61E-02 | ↑ |
| F1MAA7 | Laminin subunit gamma 1 | 2.177 | 1.40E-02 | ↑ |
| A0A0G2JV12 | Collagen type XV alpha 1 chain | 2.241 | 1.76E-02 | ↑ |
| A0A0G2K0Q8 | Sushi domain containing 2 | 2.251 | 1.66E-02 | ↑ |
| P16975 | SPARC | 2.328 | 2.02E-02 | ↑ |
| A0A0G2JY48 | receptor protein-tyrosine kinase | 2.334 | 1.67E-03 | ↑ |
| Q3MHS9 | T-complex protein 1 subunit zeta | 2.336 | 3.94E-02 | ↑ |
| E9PSU8 | Ig-like domain-containing protein | 2.338 | 1.73E-02 | ↑ |
| P61972 | Nuclear transport factor 2 | 2.346 | 2.42E-02 | ↑ |
| Q09326 | Alpha-1,6-mannosyl-glycoprotein 2-beta-N-acetylglucosaminyltransferase | 2.405 | 3.67E-02 | ↑ |
| B2RYC9 | Glucosylceramidase | 2.417 | 2.38E-02 | ↑ |
| A0A140TAI2 | Biotinidase | 2.447 | 6.64E-03 | ↑ |
| F1LSV0 | Semaphorin 4B | 2.478 | 3.81E-02 | ↑ |
| F1LMM4 | Protein kinase domain containing, cytoplasmic | 2.481 | 4.74E-02 | ↑ |
| G3V6H1 | Galactocerebrosidase | 2.490 | 1.41E-02 | ↑ |
| D3ZBN3 | receptor protein-tyrosine kinase | 2.496 | 3.87E-02 | ↑ |
| P27867 | Sorbitol dehydrogenase | 2.541 | 2.72E-02 | ↑ |
| G3V7W1 | Programmed cell death protein 6 | 2.595 | 2.93E-02 | ↑ |
| P00731 | Carboxypeptidase A1 | 2.686 | 4.49E-02 | ↑ |
| P05197 | Elongation factor 2 | 2.839 | 3.13E-02 | ↑ |
| B1WBV2 | Secreted frizzled-related protein 2 | 2.917 | 1.49E-02 | ↑ |
| Q5U367 | Multifunctional procollagen lysine hydroxylase and glycosyltransferase LH3 [Includes: Procollagen-lysine,2-oxoglutarate 5-dioxygenase 3 | 2.983 | 2.14E-02 | ↑ |
| D3ZAN3 | Glucosidase II alpha subunit | 3.074 | 1.21E-02 | ↑ |
| D3ZBB2 | deleted | 3.255 | 1.13E-02 | ↑ |
| P45479 | Palmitoyl-protein thioesterase 1 | 3.266 | 1.42E-02 | ↑ |
| O35142 | Coatomer subunit beta' | 3.325 | 2.60E-02 | ↑ |
| B5DEI2 | Amine oxidase | 3.407 | 1.62E-02 | ↑ |
| B1H232 | Immunoglobulin superfamily containing leucine-rich repeat | 3.410 | 8.35E-03 | ↑ |
| P04905 | Glutathione S-transferase Mu 1 | 3.442 | 3.66E-02 | ↑ |
| Q562C9 | Acireductone dioxygenase | 3.475 | 4.93E-03 | ↑ |
| Q80WD1 | Reticulon-4 receptor-like 2 | 3.557 | 4.87E-03 | ↑ |
| D3ZE93 | CEA cell adhesion molecule 19 | 3.606 | 1.88E-02 | ↑ |
| Q920A6 | Retinoid-inducible serine carboxypeptidase | 3.715 | 3.39E-03 | ↑ |
| Q66H12 | Alpha-N-acetylgalactosaminidase | 3.730 | 4.56E-03 | ↑ |
| F1LP42 | Hedgehog protein | 4.034 | 4.13E-03 | ↑ |
| A0A0H2UHH2 | Amyloid P component, serum | 4.067 | 1.05E-03 | ↑ |
| G3V827 | Kynurenine aminotransferase 1 | 4.172 | 4.31E-02 | ↑ |
| Q9Z339 | Glutathione S-transferase omega-1 | 4.176 | 1.84E-02 | ↑ |
| Q4FZV0 | Beta-mannosidase | 4.251 | 9.96E-03 | ↑ |
| O35760 | Isopentenyl-diphosphate Delta-isomerase 1 | 4.294 | 2.14E-02 | ↑ |
| A2IBE0 | Membrane-bound carbonic anhydrase 14 | 4.436 | 8.45E-03 | ↑ |
| M0RAM5 | Glutathione peroxidase | 4.455 | 3.51E-02 | ↑ |
| Q9EQV6 | Tripeptidyl-peptidase 1 | 4.803 | 4.37E-04 | ↑ |
| Q8R5M3 | Leucine-rich repeat-containing protein 15 | 5.157 | 4.21E-04 | ↑ |
| P84039 | Ectonucleotide pyrophosphatase/phosphodiesterase family member 5 | 5.291 | 4.31E-03 | ↑ |
| Q99M75 | Reticulon-4 receptor | 5.372 | 3.36E-02 | ↑ |
| F1M7B3 | Ig-like domain-containing protein | 5.421 | 7.84E-03 | ↑ |
| G3V862 | Angiopoietin-like 2 | 5.789 | 8.65E-03 | ↑ |
| Q5PPH0 | Enolase-phosphatase E1 | 5.794 | 4.31E-02 | ↑ |
| Q32KJ6 | N-acetylgalactosamine-6-sulfatase | 5.809 | 2.21E-02 | ↑ |
| Q5I0D5 | Phospholysine phosphohistidine inorganic pyrophosphate phosphatase | 6.164 | 1.55E-02 | ↑ |
| D3ZXJ0 | Matrix metallopeptidase 17 | 6.260 | 4.42E-02 | ↑ |
| B2GUX7 | Cellular repressor of E1A-stimulated genes 1 | 6.517 | 3.59E-02 | ↑ |
| F1M3Y4 | RCG64257-like | 7.480 | 4.96E-02 | ↑ |
| A0A0G2K5X3 | deleted | 7.517 | 3.25E-02 | ↑ |
| Q5FVH2 | 5'-3' exonuclease PLD3 | 7.683 | 1.16E-02 | ↑ |
| P53369 | Oxidized purine nucleoside triphosphate hydrolase | 9.404 | 5.06E-03 | ↑ |
| D3ZP44 | SLIT and NTRK-like family, member 6 | 10.530 | 4.31E-02 | ↑ |
| F1M091 | Kallikrein m | 15.790 | 4.14E-02 | ↑ |
| Q99ML5 | Prenylcysteine oxidase 1 | 17.347 | 4.97E-02 | ↑ |
| Q5M819 | Phosphoserine phosphatase | 34.176 | 2.31E-02 | ↑ |
| P48032 | Metalloproteinase inhibitor 3 | 306.177 | 1.05E-02 | ↑ |

Appendix Table 3 Differentially Expressed Proteins in the Urinary Proteome of the Lard Group and the Control Group Screened with FC≥1.5 or ≤0.67, P < 0.05

| Uniprot ID | Protein name | Fold change | P-value | Trend |
| --- | --- | --- | --- | --- |
| P21708 | Mitogen-activated protein kinase 3 | 0.000 | 2.67E-02 | ↓ |
| A0A0G2K1V9 | deleted | 0.017 | 2.51E-02 | ↓ |
| Q66H69 | UDP-GlcNAc:betaGal beta-1,3-N-acetylglucosaminyltransferase 7 | 0.020 | 4.93E-02 | ↓ |
| F1LMP1 | Alpha-1,3-mannosyl-glycoprotein 4-beta-N-acetylglucosaminyltransferase A | 0.036 | 3.30E-02 | ↓ |
| F1M3T3 | deleted | 0.059 | 2.98E-02 | ↓ |
| P18614 | Integrin alpha-1 | 0.080 | 3.00E-02 | ↓ |
| P35467 | Protein S100-A1 | 0.111 | 2.99E-02 | ↓ |
| B1H219 | Dickkopf WNT signaling pathway inhibitor 3 | 0.120 | 4.74E-02 | ↓ |
| Q63434 | Placenta growth factor | 0.158 | 4.47E-02 | ↓ |
| D3ZPM7 | ADAM metallopeptidase domain 19 | 0.160 | 2.52E-02 | ↓ |
| A0A0G2JXY4 | Carbonic anhydrase 15 | 0.160 | 2.14E-02 | ↓ |
| P14844 | C-C motif chemokine 2 | 0.174 | 2.49E-02 | ↓ |
| Q68FR6 | Elongation factor 1-gamma | 0.175 | 2.04E-02 | ↓ |
| Q5U206 | Calmodulin-like protein 3 | 0.298 | 4.67E-02 | ↓ |
| B4F7A5 | Cd99 protein | 0.345 | 3.26E-02 | ↓ |
| Q9JIK1 | Cadherin-related family member 5 | 0.385 | 2.81E-02 | ↓ |
| P34058 | Heat shock protein HSP 90-beta | 0.425 | 3.00E-02 | ↓ |
| A0A0G2JXZ9 | protein-tyrosine-phosphatase | 0.427 | 1.75E-02 | ↓ |
| A0A0G2K872 | Cell adhesion molecule 3 | 0.451 | 2.82E-02 | ↓ |
| Q9EPF2 | Cell surface glycoprotein MUC18 | 0.498 | 4.69E-02 | ↓ |
| P25236 | Selenoprotein P | 0.515 | 3.25E-02 | ↓ |
| D3Z9U8 | S100 calcium binding protein A7 like 2 | 0.525 | 3.42E-02 | ↓ |
| G3V6A0 | Platelet-derived growth factor receptor alpha | 0.538 | 4.91E-02 | ↓ |
| D4A6K7 | Leucine rich repeat containing 19 | 0.580 | 2.03E-02 | ↓ |
| P08289 | Alkaline phosphatase, tissue-nonspecific isozyme | 0.617 | 4.42E-02 | ↓ |
| B2RYB8 | Integrin beta | 1.541 | 9.73E-03 | ↑ |
| Q9WVH8 | Fibulin-5 | 1.584 | 4.17E-02 | ↑ |
| G3V7L8 | V-type proton ATPase subunit E 1 | 1.681 | 4.81E-03 | ↑ |
| A0A0G2KB31 | Hepsin | 1.734 | 3.77E-03 | ↑ |
| P25113 | Phosphoglycerate mutase 1 | 1.862 | 9.98E-03 | ↑ |
| M0R3V4 | Myeloid-derived growth factor | 1.916 | 4.70E-02 | ↑ |
| P80254 | D-dopachrome decarboxylase | 1.929 | 1.74E-02 | ↑ |
| P85971 | 6-phosphogluconolactonase | 1.997 | 3.14E-02 | ↑ |
| D4AC39 | Fibronectin leucine rich transmembrane protein 1 | 2.016 | 3.09E-02 | ↑ |
| Q63530 | Phosphotriesterase-related protein | 2.025 | 1.44E-02 | ↑ |
| F1LSV0 | Semaphorin 4B | 2.032 | 1.85E-02 | ↑ |
| A0A0G2JU82 | Microtubule-actin crosslinking factor 1 | 2.102 | 2.80E-02 | ↑ |
| Q711G3 | Isoamyl acetate-hydrolyzing esterase 1 homolog | 2.103 | 1.19E-02 | ↑ |
| A0A140TAI2 | Biotinidase | 2.114 | 1.04E-02 | ↑ |
| Q5XI77 | Annexin | 2.204 | 1.63E-02 | ↑ |
| P13265 | Glypican-3 | 2.230 | 2.23E-02 | ↑ |
| P19804 | Nucleoside diphosphate kinase B | 2.281 | 1.78E-02 | ↑ |
| P63322 | Ras-related protein Ral-A | 2.289 | 7.84E-03 | ↑ |
| Q32KK2 | Arylsulfatase A | 2.304 | 4.07E-03 | ↑ |
| O35142 | Coatomer subunit beta' | 2.320 | 4.96E-02 | ↑ |
| A0A0G2JSJ8 | Alpha-L-fucosidase | 2.376 | 1.81E-02 | ↑ |
| F1M8B7 | Charged multivesicular body protein 2B | 2.480 | 9.79E-03 | ↑ |
| P61972 | Nuclear transport factor 2 | 2.494 | 2.47E-04 | ↑ |
| M0RAM5 | Glutathione peroxidase | 2.506 | 2.23E-03 | ↑ |
| Q9Z339 | Glutathione S-transferase omega-1 | 2.522 | 3.47E-02 | ↑ |
| Q8CGS4 | Charged multivesicular body protein 3 | 2.538 | 1.74E-02 | ↑ |
| Q3MIE4 | Synaptic vesicle membrane protein VAT-1 homolog | 2.538 | 2.27E-02 | ↑ |
| Q5BK81 | Prostaglandin reductase 2 | 2.557 | 2.02E-02 | ↑ |
| O35760 | Isopentenyl-diphosphate Delta-isomerase 1 | 2.680 | 3.10E-02 | ↑ |
| Q6IMK4 | Amine oxidase | 2.722 | 4.54E-02 | ↑ |
| G3V7W1 | Programmed cell death protein 6 | 2.745 | 2.27E-03 | ↑ |
| G3V6H1 | Galactocerebrosidase | 2.852 | 1.20E-02 | ↑ |
| Q5XI89 | NXPE family member 4 | 2.876 | 8.73E-03 | ↑ |
| G3V8T4 | DNA damage-binding protein 1 | 2.878 | 2.68E-02 | ↑ |
| G3V8P3 | Cadherin, EGF LAG seven-pass G-type receptor 2 | 2.891 | 2.80E-02 | ↑ |
| G3V722 | Beta-1,4-galactosyltransferase | 2.921 | 3.98E-02 | ↑ |
| Q5U367 | Multifunctional procollagen lysine hydroxylase and glycosyltransferase LH3 [Includes: Procollagen-lysine,2-oxoglutarate 5-dioxygenase 3 | 2.973 | 2.87E-03 | ↑ |
| B1H232 | Immunoglobulin superfamily containing leucine-rich repeat | 2.976 | 1.25E-03 | ↑ |
| P08010 | Glutathione S-transferase Mu 2 | 3.188 | 2.56E-02 | ↑ |
| B0BNL6 | Arrestin domain-containing protein 1 | 3.191 | 2.27E-02 | ↑ |
| P02651 | Apolipoprotein A-IV | 3.246 | 4.63E-02 | ↑ |
| Q99M75 | Reticulon-4 receptor | 3.285 | 9.57E-03 | ↑ |
| D3Z9E1 | Elastin microfibril interfacer 1 | 3.338 | 2.03E-02 | ↑ |
| P08753 | Guanine nucleotide-binding protein G | 3.354 | 3.85E-02 | ↑ |
| Q66H12 | Alpha-N-acetylgalactosaminidase | 3.454 | 2.90E-02 | ↑ |
| P84039 | Ectonucleotide pyrophosphatase/phosphodiesterase family member 5 | 3.457 | 3.43E-02 | ↑ |
| Q64057 | Alpha-aminoadipic semialdehyde dehydrogenase | 3.567 | 2.53E-02 | ↑ |
| A0A0H2UHH2 | Amyloid P component, serum | 3.582 | 1.07E-03 | ↑ |
| Q99MF4 | Interleukin-11 receptor subunit alpha | 3.611 | 9.95E-03 | ↑ |
| Q920A6 | Retinoid-inducible serine carboxypeptidase | 3.837 | 2.78E-02 | ↑ |
| G3V7N8 | Galectin 9 | 3.996 | 4.84E-02 | ↑ |
| Q32KJ6 | N-acetylgalactosamine-6-sulfatase | 3.997 | 2.82E-02 | ↑ |
| P22734 | Catechol O-methyltransferase | 4.360 | 2.81E-02 | ↑ |
| A0A0G2K828 | Ig-like domain-containing protein | 4.406 | 3.31E-02 | ↑ |
| B2GUX7 | Cellular repressor of E1A-stimulated genes 1 | 4.482 | 2.15E-03 | ↑ |
| D3ZC04 | Extended synaptotagmin 3 | 4.681 | 4.39E-03 | ↑ |
| Q8R5M3 | Leucine-rich repeat-containing protein 15 | 4.748 | 3.96E-02 | ↑ |
| A2IBE0 | Membrane-bound carbonic anhydrase 14 | 4.763 | 1.32E-02 | ↑ |
| D4ACR1 | RCG64219-like | 5.020 | 3.70E-02 | ↑ |
| Q5I0D5 | Phospholysine phosphohistidine inorganic pyrophosphate phosphatase | 5.063 | 1.30E-02 | ↑ |
| A0A0G2KAY8 | Selenoprotein F | 5.094 | 2.71E-02 | ↑ |
| Q9EQV6 | Tripeptidyl-peptidase 1 | 5.099 | 6.49E-04 | ↑ |
| A0A0G2K4B4 | acid phosphatase | 5.532 | 4.62E-02 | ↑ |
| Q5PPH0 | Enolase-phosphatase E1 | 5.755 | 2.11E-02 | ↑ |
| D4A4U3 | Magnesium-dependent phosphatase 1 | 6.335 | 1.78E-02 | ↑ |
| Q6IE17 | Stefin A2-like 1 | 6.491 | 4.71E-02 | ↑ |
| Q5FVH2 | 5'-3' exonuclease PLD3 | 6.568 | 2.80E-03 | ↑ |
| P06214 | Delta-aminolevulinic acid dehydratase | 7.207 | 1.92E-02 | ↑ |
| Q8R5M5 | 2-amino-3-carboxymuconate-6-semialdehyde decarboxylase | 7.322 | 1.31E-03 | ↑ |
| A0A0G2JSZ5 | Protein disulfide-isomerase A6 | 8.080 | 4.80E-02 | ↑ |
| Q8CJ52 | Prominin-2 | 8.125 | 4.53E-02 | ↑ |
| D3ZTV3 | Leucine-rich repeat transmembrane protein FLRT2 | 8.840 | 3.17E-02 | ↑ |
| P53369 | Oxidized purine nucleoside triphosphate hydrolase | 9.126 | 5.87E-04 | ↑ |
| B2GV72 | Carbonyl reductase [NADPH] 3 | 11.616 | 8.39E-04 | ↑ |
| P03994 | Hyaluronan and proteoglycan link protein 1 | 13.005 | 5.51E-03 | ↑ |
| A0A0G2K0W3 | Protein tweety homolog | 13.487 | 1.93E-02 | ↑ |
| Q5M819 | Phosphoserine phosphatase | 25.782 | 8.88E-03 | ↑ |
| P02696 | Retinol-binding protein 1 | 38.811 | 3.04E-02 | ↑ |
| P81556 | Metalloproteinase inhibitor 4 | 85.055 | 3.60E-02 | ↑ |

Appendix Table 4 Differentially Expressed Proteins in the Urinary Proteome of the Hydrogenated Vegetable Oil Group and the Control Group Screened with FC≥1.5 or ≤0.67, P < 0.05

| Uniprot ID | Protein name | Fold change | P-value | Trend |
| --- | --- | --- | --- | --- |
| F1LMP9 | Disabled homolog 2 | 0.000 | 4.72E-02 | ↓ |
| Q5U206 | Calmodulin-like protein 3 | 0.011 | 4.55E-02 | ↓ |
| P18614 | Integrin alpha-1 | 0.036 | 2.67E-02 | ↓ |
| F1LMP1 | Alpha-1,3-mannosyl-glycoprotein 4-beta-N-acetylglucosaminyltransferase A | 0.041 | 4.72E-02 | ↓ |
| F1M3T3 | deleted | 0.074 | 3.16E-02 | ↓ |
| Q5XIF6 | Tubulin alpha-4A chain | 0.087 | 4.61E-02 | ↓ |
| A0A0G2K0V8 | Nuclear distribution C, dynein complex regulator | 0.091 | 3.39E-02 | ↓ |
| P81155 | Voltage-dependent anion-selective channel protein 2 | 0.092 | 2.31E-02 | ↓ |
| F1LM19 | Alpha-2-HS-glycoprotein | 0.116 | 4.48E-02 | ↓ |
| Q62662 | Tyrosine-protein kinase FRK | 0.142 | 4.75E-02 | ↓ |
| D3ZV56 | Ring finger protein 150 | 0.165 | 4.24E-02 | ↓ |
| P14844 | C-C motif chemokine 2 | 0.186 | 2.09E-02 | ↓ |
| Q8K4G9 | Podocin | 0.187 | 4.54E-02 | ↓ |
| D4A2Z2 | Eppin | 0.192 | 2.36E-02 | ↓ |
| Q62975 | Protein Z-dependent protease inhibitor | 0.197 | 3.62E-02 | ↓ |
| Q8R2H2 | Integrin beta-3 | 0.287 | 2.57E-02 | ↓ |
| Q63434 | Placenta growth factor | 0.306 | 2.35E-02 | ↓ |
| P41498 | Low molecular weight phosphotyrosine protein phosphatase | 0.320 | 4.89E-02 | ↓ |
| A0A0G2K2L1 | deleted | 0.329 | 3.89E-02 | ↓ |
| P16296 | Coagulation factor IX | 0.357 | 3.56E-02 | ↓ |
| F1M8E9 | lysozyme | 0.368 | 2.26E-02 | ↓ |
| Q63768 | Adapter molecule crk | 0.407 | 3.13E-02 | ↓ |
| Q3T1J1 | Eukaryotic translation initiation factor 5A-1 | 0.417 | 4.90E-02 | ↓ |
| A0A0G2K7Y0 | CD46 molecule | 0.429 | 3.01E-02 | ↓ |
| Q9QYL8 | Acyl-protein thioesterase 2 | 0.438 | 5.51E-03 | ↓ |
| Q99PW7 | Follistatin-related protein 3 | 0.441 | 1.67E-02 | ↓ |
| A0A0G2K676 | chitinase | 0.442 | 2.87E-02 | ↓ |
| P02680 | Fibrinogen gamma chain | 0.475 | 4.45E-02 | ↓ |
| F1LQT4 | Carboxypeptidase N subunit 2 | 0.480 | 2.41E-02 | ↓ |
| Q9EPF2 | Cell surface glycoprotein MUC18 | 0.522 | 3.91E-02 | ↓ |
| G3V7Y3 | ATP synthase F1 subunit delta | 0.533 | 3.87E-02 | ↓ |
| D3ZCH9 | CD177 molecule | 0.541 | 4.60E-02 | ↓ |
| Q99J86 | Attractin | 0.577 | 6.75E-03 | ↓ |
| G3V6K1 | Transcobalamin-2 | 0.591 | 2.26E-02 | ↓ |
| Q9QX71 | Napsin | 1.549 | 1.57E-02 | ↑ |
| P85971 | 6-phosphogluconolactonase | 1.640 | 3.52E-02 | ↑ |
| Q80ZA3 | Serine | 1.699 | 4.24E-02 | ↑ |
| F1M1R0 | Similar to immunoglobulin light chain variable region | 1.788 | 3.80E-02 | ↑ |
| F1LSV0 | Semaphorin 4B | 1.847 | 3.57E-02 | ↑ |
| B2RYC9 | Glucosylceramidase | 1.865 | 3.69E-02 | ↑ |
| D3ZC04 | Extended synaptotagmin 3 | 2.014 | 3.45E-02 | ↑ |
| P02651 | Apolipoprotein A-IV | 2.120 | 2.91E-02 | ↑ |
| A0A0H2UHQ0 | Solute carrier family 3 member 2 | 2.145 | 3.79E-02 | ↑ |
| Q562C9 | Acireductone dioxygenase | 2.276 | 2.49E-02 | ↑ |
| Q80WD1 | Reticulon-4 receptor-like 2 | 2.365 | 3.08E-02 | ↑ |
| D3ZAN3 | Glucosidase II alpha subunit | 2.365 | 2.30E-02 | ↑ |
| Q99MF4 | Interleukin-11 receptor subunit alpha | 2.603 | 2.19E-02 | ↑ |
| Q64573 | Liver carboxylesterase 1F | 2.625 | 4.17E-02 | ↑ |
| G3V862 | Angiopoietin-like 2 | 2.626 | 3.13E-02 | ↑ |
| O35760 | Isopentenyl-diphosphate Delta-isomerase 1 | 2.763 | 3.37E-02 | ↑ |
| Q64361 | Latexin | 2.916 | 3.41E-02 | ↑ |
| A0A0G2K0Q8 | Sushi domain containing 2 | 3.023 | 3.66E-02 | ↑ |
| A0A0H2UHH2 | Amyloid P component, serum | 3.156 | 3.78E-02 | ↑ |
| D4AA31 | Prolylcarboxypeptidase | 3.491 | 3.78E-02 | ↑ |
| B0BN46 | Grhpr protein | 3.681 | 1.92E-02 | ↑ |
| Q9EQV6 | Tripeptidyl-peptidase 1 | 3.706 | 4.88E-02 | ↑ |
| E9PSU8 | Ig-like domain-containing protein | 3.714 | 3.60E-02 | ↑ |
| P22734 | Catechol O-methyltransferase | 3.841 | 2.40E-02 | ↑ |
| M0R936 | Ig-like domain-containing protein | 4.155 | 4.81E-02 | ↑ |
| Q5FVH2 | 5'-3' exonuclease PLD3 | 4.715 | 2.20E-02 | ↑ |
| D4A4U3 | Magnesium-dependent phosphatase 1 | 5.130 | 2.60E-02 | ↑ |
| Q8R5M5 | 2-amino-3-carboxymuconate-6-semialdehyde decarboxylase | 10.148 | 2.71E-02 | ↑ |
| Q5M819 | Phosphoserine phosphatase | 14.636 | 4.44E-02 | ↑ |

Appendix Table 5 Differentially Expressed Proteins in the Urinary Proteome of the Rapeseed Oil Group and the Control Group Screened with FC≥1.5 or ≤0.67, P < 0.05

| Uniprot ID | Protein name | Fold change | P-value | Trend |
| --- | --- | --- | --- | --- |
| A0A096MIU6 | Fc fragment of IgG receptor IIa | 0.000 | 4.14E-02 | ↓ |
| A0A0G2JTL4 | Mast/stem cell growth factor receptor | 0.000 | 3.33E-02 | ↓ |
| A0A0G2K0V8 | Nuclear distribution C, dynein complex regulator | 0.000 | 2.81E-02 | ↓ |
| A0A0G2K7L8 | Thrombospondin 4 | 0.000 | 4.46E-02 | ↓ |
| D3ZM36 | Interleukin 10 receptor subunit beta | 0.000 | 2.63E-02 | ↓ |
| D3ZV56 | Ring finger protein 150 | 0.000 | 2.83E-02 | ↓ |
| D3ZW55 | Inosine triphosphate pyrophosphatase | 0.000 | 4.44E-02 | ↓ |
| D4A740 | Interleukin 17 receptor A | 0.000 | 2.50E-02 | ↓ |
| D4ACX8 | Protocadherin-16 | 0.000 | 4.93E-02 | ↓ |
| F1LMP9 | Disabled homolog 2 | 0.000 | 4.72E-02 | ↓ |
| F1LW74 | IQ motif containing GTPase activating protein 2 | 0.000 | 2.72E-02 | ↓ |
| F1M7W8 | deleted | 0.000 | 2.65E-02 | ↓ |
| P16975 | SPARC | 0.000 | 4.05E-02 | ↓ |
| P18614 | Integrin alpha-1 | 0.000 | 2.42E-02 | ↓ |
| P35467 | Protein S100-A1 | 0.000 | 3.86E-02 | ↓ |
| Q4V8I1 | Endothelial protein C receptor | 0.000 | 3.81E-02 | ↓ |
| Q5XIF6 | Tubulin alpha-4A chain | 0.000 | 3.68E-02 | ↓ |
| Q5XIN7 | Protein O-linked-mannose beta-1,2-N-acetylglucosaminyltransferase 1 | 0.000 | 3.26E-02 | ↓ |
| Q66H69 | UDP-GlcNAc:betaGal beta-1,3-N-acetylglucosaminyltransferase 7 | 0.000 | 4.89E-02 | ↓ |
| Q9ES58 | Cell surface glycoprotein CD200 receptor 1 | 0.000 | 4.12E-02 | ↓ |
| Q63434 | Placenta growth factor | 0.000 | 3.27E-02 | ↓ |
| Q5U206 | Calmodulin-like protein 3 | 0.002 | 4.41E-02 | ↓ |
| F1LMP1 | Alpha-1,3-mannosyl-glycoprotein 4-beta-N-acetylglucosaminyltransferase A | 0.016 | 3.58E-02 | ↓ |
| A0A0G2JVV1 | C-type lectin domain family 2, member G | 0.040 | 1.77E-02 | ↓ |
| P70549 | Sodium/calcium exchanger 3 | 0.044 | 4.00E-02 | ↓ |
| Q9ES53 | Ubiquitin recognition factor in ER-associated degradation protein 1 | 0.047 | 2.33E-02 | ↓ |
| D3ZPM7 | ADAM metallopeptidase domain 19 | 0.047 | 2.18E-02 | ↓ |
| Q5I0D1 | Glyoxalase domain-containing protein 4 | 0.050 | 4.90E-02 | ↓ |
| Q62975 | Protein Z-dependent protease inhibitor | 0.057 | 2.61E-02 | ↓ |
| A0A0G2KA95 | L1 cell adhesion molecule | 0.059 | 2.42E-02 | ↓ |
| A0A0G2JZ40 | Reversion-inducing-cysteine-rich protein with kazal motifs | 0.062 | 3.54E-02 | ↓ |
| M0R7T1 | Phosphatase and actin regulator 4 | 0.066 | 1.38E-02 | ↓ |
| D4ACM8 | Frizzled class receptor 7 | 0.070 | 3.97E-02 | ↓ |
| Q68HB8 | Protocadherin 7 | 0.072 | 4.34E-02 | ↓ |
| P32736 | Opioid-binding protein/cell adhesion molecule | 0.073 | 4.90E-02 | ↓ |
| P06399 | Fibrinogen alpha chain [Cleaved into: Fibrinopeptide A; Fibrinogen alpha chain] | 0.077 | 2.83E-02 | ↓ |
| A0A0G2JXG3 | Cell cycle control protein | 0.084 | 4.05E-02 | ↓ |
| A0A0G2JY19 | Hepatocyte growth factor receptor | 0.098 | 1.78E-02 | ↓ |
| Q499S6 | Cathepsin F | 0.142 | 2.35E-02 | ↓ |
| A0A0G2JXP0 | deleted | 0.146 | 1.89E-02 | ↓ |
| P13803 | Electron transfer flavoprotein subunit alpha, mitochondrial | 0.168 | 2.84E-02 | ↓ |
| P02680 | Fibrinogen gamma chain | 0.170 | 2.96E-02 | ↓ |
| G3V8N9 | Testis expressed 101 | 0.225 | 2.27E-02 | ↓ |
| A0A0G2JXY4 | Carbonic anhydrase 15 | 0.262 | 1.18E-02 | ↓ |
| A0A0G2K994 | V-set and immunoglobulin domain containing 10 | 0.283 | 2.07E-02 | ↓ |
| Q8R2H2 | Integrin beta-3 | 0.283 | 3.88E-02 | ↓ |
| Q9QYL8 | Acyl-protein thioesterase 2 | 0.285 | 7.95E-03 | ↓ |
| Q6AYK3 | Inositol-3-phosphate synthase 1 | 0.327 | 2.30E-02 | ↓ |
| P25236 | Selenoprotein P | 0.346 | 3.66E-03 | ↓ |
| Q9JIK1 | Cadherin-related family member 5 | 0.365 | 4.87E-02 | ↓ |
| A0A0G2JXZ9 | protein-tyrosine-phosphatase | 0.379 | 4.83E-02 | ↓ |
| P34058 | Heat shock protein HSP 90-beta | 0.405 | 3.25E-04 | ↓ |
| B1WBS5 | Sodium/mannose cotransporter SLC5A10 | 0.453 | 4.40E-02 | ↓ |
| A0A0G2K980 | Ig-like domain-containing protein | 1.661 | 3.07E-02 | ↑ |
| M0R7Q2 | Ig-like domain-containing protein | 1.693 | 3.52E-02 | ↑ |
| M0RC20 | deleted | 1.873 | 2.15E-02 | ↑ |
| P61972 | Nuclear transport factor 2 | 1.986 | 1.27E-02 | ↑ |
| D3ZJF9 | Alpha-galactosidase | 2.055 | 1.70E-02 | ↑ |
| A0A0G2JXI1 | Ac2-120 | 2.057 | 1.11E-02 | ↑ |
| P47967 | Galectin-5 | 2.145 | 1.92E-02 | ↑ |
| Z4YNX7 | Cystatin-related protein 2 | 2.149 | 2.02E-02 | ↑ |
| P10758 | Lithostathine | 2.235 | 3.29E-02 | ↑ |
| Q5I0E1 | Leucine-rich alpha-2-glycoprotein 1 | 2.333 | 1.41E-03 | ↑ |
| M0R7M5 | deleted | 2.410 | 3.83E-02 | ↑ |
| P70490 | Lactadherin | 2.753 | 3.89E-02 | ↑ |
| A0A0G2K3W2 | Coagulation factor V | 2.791 | 2.80E-02 | ↑ |
| A0A140TAI2 | Biotinidase | 2.842 | 8.33E-03 | ↑ |
| Q80ZA3 | Serine peptidase inhibitor, clade F, member 1 | 3.127 | 1.30E-02 | ↑ |
| Q6AXR4 | Beta-hexosaminidase subunit beta | 3.177 | 1.46E-02 | ↑ |
| A0A0H2UHH2 | Amyloid P component, serum | 3.260 | 2.58E-02 | ↑ |
| P30152 | Neutrophil gelatinase-associated lipocalin | 3.267 | 1.91E-02 | ↑ |
| Q00657 | Chondroitin sulfate proteoglycan 4 | 3.297 | 2.06E-02 | ↑ |
| Q5XI77 | Annexin | 3.346 | 8.95E-03 | ↑ |
| M0RDA4 | receptor protein-tyrosine kinase | 3.503 | 1.40E-02 | ↑ |
| A0A0G2JY48 | receptor protein-tyrosine kinase | 3.504 | 5.89E-03 | ↑ |
| A0A0G2K5X3 | deleted | 3.533 | 3.88E-02 | ↑ |
| Q641X3 | Beta-hexosaminidase subunit alpha | 3.549 | 4.40E-02 | ↑ |
| Q9QZK9 | Deoxyribonuclease-2-beta | 3.555 | 4.08E-02 | ↑ |
| Q920A6 | Retinoid-inducible serine carboxypeptidase | 3.638 | 4.99E-03 | ↑ |
| D4A4R7 | Serpin family A member 1F | 3.757 | 1.76E-02 | ↑ |
| F1MA98 | Nucleoprotein TPR | 4.262 | 3.22E-02 | ↑ |
| P45479 | Palmitoyl-protein thioesterase 1 | 4.525 | 3.34E-02 | ↑ |
| Q5I0D5 | Phospholysine phosphohistidine inorganic pyrophosphate phosphatase | 4.774 | 2.79E-02 | ↑ |
| G3V7D0 | Matrix metallopeptidase 8 | 4.890 | 1.98E-02 | ↑ |
| A0A0G2JZN1 | deleted | 5.142 | 4.03E-02 | ↑ |
| Q32KK2 | Arylsulfatase A | 5.412 | 2.33E-02 | ↑ |
| G3V862 | Angiopoietin-like 2 | 5.826 | 3.90E-02 | ↑ |
| Q9Z2Y9 | Klotho | 6.035 | 1.23E-02 | ↑ |
| Q32KJ6 | N-acetylgalactosamine-6-sulfatase | 6.364 | 3.41E-02 | ↑ |
| B2RYC9 | Glucosylceramidase | 6.461 | 5.14E-03 | ↑ |
| Q80WD1 | Reticulon-4 receptor-like 2 | 6.905 | 4.90E-02 | ↑ |
| Q6P7S1 | Acid ceramidase | 7.057 | 2.43E-02 | ↑ |
| D3ZY02 | Protein-glucosylgalactosylhydroxylysine glucosidase | 7.293 | 2.83E-02 | ↑ |
| F1MAN8 | Laminin subunit alpha 5 | 7.979 | 4.31E-02 | ↑ |
| Q66HT5 | CCN family member 1 | 9.008 | 3.96E-02 | ↑ |
| Q6P762 | Alpha-mannosidase | 9.213 | 2.90E-02 | ↑ |
| D4ACR1 | RCG64219-like | 9.576 | 2.19E-02 | ↑ |
| D4AE30 | Prostate and testis expressed C | 9.649 | 4.20E-02 | ↑ |
| F1LXY6 | deleted | 11.107 | 1.46E-02 | ↑ |
| F1M091 | Kallikrein m | 11.174 | 3.48E-02 | ↑ |
| A0A0G2K8F6 | Alpha-mannosidase | 11.469 | 2.22E-03 | ↑ |
| G3V8X9 | Serine peptidase inhibitor | 13.180 | 4.40E-02 | ↑ |
| Q5FVH2 | 5'-3' exonuclease PLD3 | 15.910 | 3.46E-02 | ↑ |
| F1LWD0 | Ig-like domain-containing protein | 25.353 | 4.22E-02 | ↑ |
| A0A0G2K680 | Serine peptidase inhibitor, Kunitz type, 3 | 43.172 | 4.78E-02 | ↑ |

Appendix Table 6 Differential Modifications in the Urinary Proteome of the Olive Oil Group and the Control Group Screened with FC≥1.5 or ≤0.67, P < 0.05

| Uniprot ID | Peptide | Modification | A组 | B组 | Flod change | P value |
| --- | --- | --- | --- | --- | --- | --- |
| P02761 | EKIEENGSMRVFMQHIDVLENSLGFK | 0,GIST-Quat[AnyN-term]; | 2.4 | 0.0 | 0.000 | 4.18E-02 |
| P02761 | IDVLENSLGFK | 1,Dioxidation[I]; | 1.0 | 0.0 | 0.000 | 3.41E-02 |
| P0DP29 | DGNGYISAAELR | 3,Deamidated[N]; | 4.2 | 0.0 | 0.000 | 4.54E-02 |
| P0DP30 | DGNGYISAAELR | 3,Deamidated[N]; | 4.2 | 0.0 | 0.000 | 4.54E-02 |
| P0DP31 | DGNGYISAAELR | 3,Deamidated[N]; | 4.2 | 0.0 | 0.000 | 4.54E-02 |
| P14046 | KQSGVKEEHSFTVMEFVLPR | 0,SPITC_13C(6)[AnyN-term]; | 2.0 | 0.0 | 0.000 | 4.74E-02 |
| P36373 | NNLLEDEPFAQHR | 3,Xle->Gln[L]; | 1.4 | 0.0 | 0.000 | 2.49E-02 |
| P36374 | NNLLEDEPFAQHR | 3,Xle->Gln[L]; | 1.4 | 0.0 | 0.000 | 2.49E-02 |
| P97574 | MIAEVQEDCYSK | 9,Carbamidomethyl[C]; | 9.0 | 0.0 | 0.000 | 4.59E-02 |
| Q03626 | KQSGVKEEHSFTVMEFVLPR | 0,SPITC_13C(6)[AnyN-term]; | 2.0 | 0.0 | 0.000 | 4.74E-02 |
| Q6IE52 | KQSGVKEEHSFTVMEFVLPR | 0,SPITC_13C(6)[AnyN-term]; | 2.0 | 0.0 | 0.000 | 4.74E-02 |
| Q9JJ19 | IVEVNGVCMEGK | 8,Carbamidomethyl[C];9,Oxidation[M]; | 4.8 | 0.0 | 0.000 | 3.05E-02 |
| Q9JJ19 | IVEVNGVCMEGK | 8,Carbamidomethyl[C]; | 6.6 | 0.6 | 0.091 | 4.44E-02 |
| P15083 | SSVTFECDLGR | 7,Carbamidomethyl[C]; | 52.8 | 11.2 | 0.212 | 3.28E-02 |
| P81828 | LALQCFRCTSFDSTGFCHVGR | 5,Carbamidomethyl[C];17,Carbamidomethyl[C]; | 1.6 | 0.4 | 0.250 | 3.27E-02 |
| P83121 | HICQTYPDEICAWVVVTTR | 3,NEM_2H(5)[C]; | 118.4 | 35.0 | 0.296 | 4.72E-02 |
| P26644 | ITCPPPPIPK | 3,Carbamidomethyl[C]; | 27.8 | 13.6 | 0.489 | 3.10E-02 |
| P02625 | SMTDLLSAEDIKK | 0,Acetyl[ProteinN-term]; | 3.6 | 7.2 | 2.000 | 3.67E-02 |
| Q64240 | ECLQTCR | 2,Carbamidomethyl[C];6,Carbamidomethyl[C]; | 6.8 | 14.4 | 2.118 | 3.40E-02 |
| P07522 | TTTYAAAGPPR | 1,Dimethylphosphothione[T]; | 1.6 | 3.4 | 2.125 | 8.58E-03 |
| P22282 | LDNCPFEEQTEQLKR | 4,Carbamidomethyl[C]; | 70.4 | 149.6 | 2.125 | 3.04E-02 |
| P14841 | PQEADASEEGVQR | 1,Pro->Cys[P]; | 0.6 | 1.4 | 2.333 | 1.61E-02 |
| Q64240 | QGPCRAFAELWAFDAAQGK | 4,BHT[C]; | 8.8 | 21.6 | 2.455 | 4.04E-02 |
| P01835 | GVLDSVTDQDSK | 0,glycidamide[AnyN-term]; | 7.0 | 18.2 | 2.600 | 2.49E-02 |
| P01836 | GVLDSVTDQDSK | 0,glycidamide[AnyN-term]; | 7.0 | 18.2 | 2.600 | 2.49E-02 |
| P27590 | CPHTEDTTIQVTENGESSQAR | 1,Carbamidomethyl[C];14,Deamidated[N]; | 1.0 | 2.6 | 2.600 | 3.49E-02 |
| P20767 | NSFTCQVTHEGNTVEK | 1,Deamidated[N];5,Carbamidomethyl[C]; | 4.0 | 14.4 | 3.600 | 3.05E-02 |
| P01835 | DSTYSMSSTLSLTK | 6,Carbamidomethyl[M]; | 2.2 | 9.0 | 4.091 | 1.56E-04 |
| P01836 | DSTYSMSSTLSLTK | 6,Carbamidomethyl[M]; | 2.2 | 9.0 | 4.091 | 1.56E-04 |
| P98158 | CQTTNICVPR | 1,Carbamidomethyl[C];7,Carbamidomethyl[C]; | 1.0 | 4.6 | 4.600 | 3.27E-02 |
| Q64230 | PVENRQAIMTILDQEPDAR | 0,SPITC_13C(6)[AnyN-term]; | 1.0 | 4.6 | 4.600 | 6.05E-03 |
| P22282 | LDNCPFEEQTEQLKR | 3,Deamidated[N];4,Carbamidomethyl[C]; | 0.6 | 2.8 | 4.667 | 4.02E-02 |
| P01835 | RDGVLDSVTDQDSK | 13,Formyl[S](Ser->Asp[S]); | 1.6 | 7.6 | 4.750 | 1.67E-02 |
| P01836 | RDGVLDSVTDQDSK | 13,Formyl[S](Ser->Asp[S]); | 1.6 | 7.6 | 4.750 | 1.67E-02 |
| P22282 | LDNCPFEEQTEQLKR | 4,CarbamidomethylDTT[C]; | 2.0 | 10.4 | 5.200 | 2.77E-02 |
| Q68FP1 | PKAGALNSNDAFVLK | 0,Unknown_420[AnyN-term]; | 1.0 | 5.8 | 5.800 | 3.27E-02 |
| P01835 | RDGVLDSVTDQDSK | 13,Carbonyl[S]; | 0.6 | 6.0 | 10.000 | 8.58E-03 |
| P01836 | RDGVLDSVTDQDSK | 13,Carbonyl[S]; | 0.6 | 6.0 | 10.000 | 8.58E-03 |
| Q6P7A9 | AVPTQCDVTPNSR | 6,Carbamidomethyl[C]; | 0.6 | 8.4 | 14.000 | 4.46E-04 |
| Q63041 | QDLNDNDAYSVFQSIGLK | 0,Propionyl_13C(3)[AnyN-term]; | 0.2 | 3.0 | 15.000 | 1.89E-02 |
| P22057 | GHDTVQPNFQQDK | 4,Thiophospho[T]; | 0.6 | 9.2 | 15.333 | 2.62E-03 |

Appendix Table 7 Differential Modifications in the Urinary Proteome of the Butter Group and the Control Group Screened with FC≥1.5 or ≤0.67, P < 0.05

| Uniprot ID | Peptide | Modification | A组 | C组 | Flod change | P value |
| --- | --- | --- | --- | --- | --- | --- |
| P30919 | YCGPYKPPDFLEQNNR | 2,Carbamidomethyl[C]; | 3.8 | 0.8 | 0.211 | 3.41E-02 |
| P02780 | SGCSILDEVIR | 3,CarbamidomethylDTT[C]; | 1.4 | 0.4 | 0.286 | 3.41E-02 |
| Q9JHB9 | SGCSILDEVIR | 3,CarbamidomethylDTT[C]; | 1.4 | 0.4 | 0.286 | 3.41E-02 |
| Q62740 | GYSVPTAACR | 9,Carbamidomethyl[C]; | 16.2 | 7.6 | 0.469 | 1.41E-02 |
| P01836 | TSSSPVVKSFNR | 8,Carbamyl[K]; | 15.0 | 24.6 | 1.640 | 4.47E-02 |
| P01835 | TSSSPVVKSFNR | 8,Carbamyl[K]; | 15.0 | 24.6 | 1.640 | 4.47E-02 |
| P22282 | LDNCPFEEQTEQLKR | 4,Carbamidomethyl[C]; | 70.4 | 128.6 | 1.827 | 6.48E-03 |
| P20767 | LTVFPPSTEELQGNK | 2,Hex(1)Pent(2)[T]; | 13.6 | 27.2 | 2.000 | 3.35E-02 |
| P20767 | NSFTCQVTHEGNTVEK | 5,Carbamidomethyl[C]; | 26.4 | 54.0 | 2.045 | 7.97E-03 |
| P22282 | LDNCPFEEQTEQLK | 4,Carbamidomethyl[C]; | 66.8 | 138.2 | 2.069 | 2.52E-03 |
| P20611 | FPLGPCPR | 6,Carbamidomethyl[C]; | 9.2 | 19.2 | 2.087 | 3.36E-02 |
| P07522 | CHELVACPGNR | 1,Carbamidomethyl[C];7,Carbamidomethyl[C]; | 2.2 | 5.2 | 2.364 | 2.31E-02 |
| P07522 | RITEGVDTPEGLAVDWIGR | 2,Xle->Asn[I]; | 1.2 | 3.2 | 2.667 | 2.17E-02 |
| P29598 | TDSCSGDSGGPLICNIDGR | 4,Carbamidomethyl[C];14,Carbamidomethyl[C]; | 5.4 | 15.4 | 2.852 | 1.59E-02 |
| P42854 | SSGNSGQNVWIGLHDPTLGQEPNR | 4,Deamidated[N];8,Deamidated[N]; | 1.0 | 3.0 | 3.000 | 2.17E-02 |
| P20767 | TLTVFPPSTEELQGNK | 1,Hex(2)[T]; | 2.8 | 9.6 | 3.429 | 1.30E-02 |
| P02761 | WFSIVVASNK | 3,Hex(1)Pent(2)[S]; | 3.0 | 10.6 | 3.533 | 4.40E-02 |
| P09006 | SMEEILEGLK | 0,Biotin[AnyN-term]; | 1.6 | 6.0 | 3.750 | 4.85E-02 |
| P07522 | VNLDPASVPPR | 2,Asn->Cys[N]; | 4.8 | 19.0 | 3.958 | 4.30E-02 |
| P20767 | NSFTCQVTHEGNTVEK | 1,Deamidated[N];5,Carbamidomethyl[C]; | 4.0 | 16.0 | 4.000 | 1.86E-02 |
| P42854 | IGLHDPTLGQEPNR | 4,His->Xle[H]; | 0.4 | 1.6 | 4.000 | 3.27E-02 |
| P42854 | WRENNCISELPYVCK | 4,Deamidated[N];6,Carbamidomethyl[C];14,Carbamidomethyl[C]; | 4.8 | 19.4 | 4.042 | 3.64E-02 |
| P01836 | DSTYSMSSTLSLTK | 6,Carbamidomethyl[M]; | 2.2 | 9.8 | 4.455 | 3.26E-03 |
| P01835 | DSTYSMSSTLSLTK | 6,Carbamidomethyl[M]; | 2.2 | 9.8 | 4.455 | 3.26E-03 |
| P20767 | FPPSTEELQGNK | 0,Formyl[AnyN-term]; | 0.8 | 4.0 | 5.000 | 4.01E-02 |
| P22282 | EICYFVVYPDYIEQNIHAVR | 0,Amidine[AnyN-term]; | 0.2 | 1.0 | 5.000 | 1.61E-02 |
| P36373 | KPGDDHSNDLMLLHLSQPADITDGVK | 5,Cation_Fe[III][D]; | 0.4 | 2.0 | 5.000 | 3.49E-02 |
| P01835 | DYESHNLYTCEVVHKTSSSPVVK | 10,Ub-VME[C]; | 0.4 | 2.4 | 6.000 | 4.74E-02 |
| P02767 | FASGKTAESGELHGLTTDEK | 6,PhosphoHexNAc[T]; | 0.4 | 2.4 | 6.000 | 4.74E-02 |
| Q4FZV0 | FQSAVQYAECQSK | 10,Carbamidomethyl[C]; | 1.0 | 6.0 | 6.000 | 3.83E-02 |
| P07522 | PSSLVVVHPLAK | 2,Dehydrated[S]; | 0.4 | 2.8 | 7.000 | 2.40E-02 |
| Q63041 | MGASEVAQEVEVR | 0,CLIP_TRAQ_2[AnyN-term]; | 0.4 | 2.8 | 7.000 | 4.18E-02 |
| Q01177 | GTPCQEWAAQEPHSHR | 4,NEM_2H(5)[C]; | 0.2 | 1.4 | 7.000 | 3.88E-03 |
| Q9EPB1 | SLPFGVQSTQR | 0,C+12[AnyN-term]; | 0.4 | 2.8 | 7.000 | 2.40E-02 |
| P22282 | PGETMYYISLPGSVR | 0,Phenylisocyanate_2H(5)[AnyN-term]; | 2.0 | 14.2 | 7.100 | 1.51E-02 |
| P20767 | QVTHEGNTVEK | 2,Val->Ser[V]; | 1.8 | 13.6 | 7.556 | 2.62E-02 |
| P02761 | LCEAHGITR | 2,maleimide[C]; | 0.6 | 5.8 | 9.667 | 3.67E-02 |
| P01835 | ADYESHNLYTCEVVHK | 7,Deamidated[N];11,Carbamidomethyl[C]; | 0.8 | 7.8 | 9.750 | 2.79E-02 |
| Q64240 | YCGVPGDGYEELTR | 2,Bromobimane[C]; | 0.4 | 4.2 | 10.500 | 3.40E-02 |
| Q6P7A9 | AVPTQCDVTPNSR | 6,Carbamidomethyl[C]; | 0.6 | 7.4 | 12.333 | 2.99E-02 |
| P01836 | RDGVLDSVTDQDSK | 14,Lys->Asp[K]; | 0.4 | 5.2 | 13.000 | 3.27E-02 |
| P01835 | RDGVLDSVTDQDSK | 14,Lys->Asp[K]; | 0.4 | 5.2 | 13.000 | 3.27E-02 |
| P01836 | RDGVLDSVTDQDSK | 13,Carbonyl[S]; | 0.6 | 8.6 | 14.333 | 3.66E-02 |
| P01835 | RDGVLDSVTDQDSK | 13,Carbonyl[S]; | 0.6 | 8.6 | 14.333 | 3.66E-02 |
| P22057 | GHDTVQPNFQQDK | 4,Thiophospho[T]; | 0.6 | 10.4 | 17.333 | 1.16E-02 |
| Q5FVF9 | NLDVYEQQVMAAAQK | 1,dHex[N]; | 0.2 | 4.8 | 24.000 | 4.78E-02 |
| P36375 | NNLFEDEPFAQYR | 0,Dimethyl_2H(4)[AnyN-term]; | 0.2 | 6.0 | 30.000 | 3.53E-02 |
| P00689 | ALVFVDNHDNQR | 6,Cation_Na[D]; | 0.2 | 6.0 | 30.000 | 3.84E-02 |
| P02770 | TVMGDFAQFVDK | 3,Met->Lys[M]; | 0.2 | 7.4 | 37.000 | 1.87E-02 |

Appendix Table 8 Differential Modifications in the Urinary Proteome of the Lard Group and the Control Group Screened with FC≥1.5 or ≤0.67, P < 0.05

| Uniprot ID | Peptide | Modification | A组 | D组 | Flod change | P value |
| --- | --- | --- | --- | --- | --- | --- |
| P02761 | EKIEENGSMRVFMQHIDVLENSLGFK | 0,GIST-Quat[AnyN-term]; | 2.4 | 0.0 | 0.000 | 4.18E-02 |
| P02761 | IDVLENSLGFK | 1,Dioxidation[I]; | 1.0 | 0.0 | 0.000 | 3.41E-02 |
| P22283 | LVNCPFEEQTEQLK | 4,dichlorination[C]; | 1.4 | 0.0 | 0.000 | 2.49E-02 |
| P32038 | DVAGWGVVTHAGR | 0,mTRAQ_13C(6)15N(2)[AnyN-term]; | 2.2 | 0.0 | 0.000 | 2.95E-02 |
| P36373 | NNLLEDEPFAQHR | 3,Xle->Gln[L]; | 1.4 | 0.0 | 0.000 | 2.49E-02 |
| P36374 | NNLLEDEPFAQHR | 3,Xle->Gln[L]; | 1.4 | 0.0 | 0.000 | 2.49E-02 |
| Q99PS8 | HPPVFGLCR | 8,Carbamidomethyl[C]; | 9.0 | 0.0 | 0.000 | 3.67E-02 |
| Q9JJ19 | IVEVNGVCMEGK | 8,Carbamidomethyl[C];9,Oxidation[M]; | 4.8 | 0.0 | 0.000 | 3.05E-02 |
| P30919 | YCGPYKPPDFLEQNNR | 2,Carbamidomethyl[C]; | 3.8 | 0.2 | 0.053 | 4.91E-02 |
| P11232 | AFQEALAAAGDK | 0,Pyridylacetyl[AnyN-term](Phenylisocyanate[AnyN-term]); | 2.2 | 0.2 | 0.091 | 3.41E-02 |
| P02761 | IDVLENSLGFK | 2,Cation_Ca[II][D]; | 4.0 | 0.4 | 0.100 | 3.27E-02 |
| P02761 | LNGDWFSIVVASNK | 4,Cation_Al[III][D]; | 8.6 | 1.0 | 0.116 | 4.78E-02 |
| P42854 | SGQNVWIGLHDPTLGQEPNR | 1,HexNAc(1)dHex(1)[S]; | 10.2 | 1.2 | 0.118 | 3.35E-02 |
| P00787 | HCGIESEIVAGIPR | 2,Cytopiloyne[C]; | 8.0 | 1.0 | 0.125 | 4.94E-02 |
| P20059 | GECQSEGVLFFQGNR | 3,Amidino[C]; | 1.6 | 0.2 | 0.125 | 2.49E-02 |
| P02780 | SGSGCSILDEVIR | 5,Malonyl[C]; | 2.4 | 0.4 | 0.167 | 3.41E-02 |
| Q9JHB9 | SGSGCSILDEVIR | 5,Malonyl[C]; | 2.4 | 0.4 | 0.167 | 3.41E-02 |
| P27590 | STDYGAGYSCDSDMHGWYR | 10,monomethylphosphothione[C]; | 2.6 | 1.0 | 0.385 | 3.49E-02 |
| P27590 | CPHTEDTTIQVTENGESSQAR | 1,CarbamidomethylDTT[C]; | 6.2 | 2.8 | 0.452 | 3.88E-02 |
| Q62740 | GYSVPTAACR | 9,Carbamidomethyl[C]; | 16.2 | 8.0 | 0.494 | 1.55E-03 |
| P81827 | ESLDSTGLCR | 9,Carbamidomethyl[C]; | 14.6 | 8.6 | 0.589 | 2.44E-02 |
| P08649 | EPFLSCCK | 6,Carbamidomethyl[C];7,Carbamidomethyl[C]; | 12.6 | 8.2 | 0.651 | 2.95E-02 |
| Q62740 | ESGDPSTCAFQR | 8,Carbamidomethyl[C]; | 19.8 | 13.0 | 0.657 | 3.64E-03 |
| P02761 | EKIEENGSMR | 9,Oxidation[M]; | 85.6 | 134.8 | 1.575 | 3.78E-02 |
| P02780 | SGSGCSILDEVIR | 5,CarbamidomethylDTT[C]; | 17.6 | 28.0 | 1.591 | 6.43E-04 |
| Q9JHB9 | SGSGCSILDEVIR | 5,CarbamidomethylDTT[C]; | 17.6 | 28.0 | 1.591 | 6.43E-04 |
| P98158 | NPCASASCSHLCLLSAQAPR | 3,Carbamidomethyl[C];8,Carbamidomethyl[C];12,Carbamidomethyl[C]; | 6.8 | 11.4 | 1.676 | 4.27E-02 |
| P98158 | HQCLCEEGYILER | 3,Carbamidomethyl[C];5,Carbamidomethyl[C]; | 18.0 | 31.6 | 1.756 | 1.97E-02 |
| P22283 | CSFQWEFCNIK | 1,Carbamidomethyl[C];8,Carbamidomethyl[C]; | 12.8 | 23.6 | 1.844 | 2.96E-02 |
| P10252 | EDQGVDYTWYEDSGPFPQR | 1,Glu->Xle[E]; | 2.8 | 5.2 | 1.857 | 9.26E-03 |
| P01835 | TSSSPVVKSFNR | 8,Carbamyl[K]; | 15.0 | 29.2 | 1.947 | 3.87E-02 |
| P01836 | TSSSPVVKSFNR | 8,Carbamyl[K]; | 15.0 | 29.2 | 1.947 | 3.87E-02 |
| P01835 | DGVLDSVTDQDSK | 1,Cation_Na[D]; | 3.6 | 7.2 | 2.000 | 2.88E-02 |
| P01836 | DGVLDSVTDQDSK | 1,Cation_Na[D]; | 3.6 | 7.2 | 2.000 | 2.88E-02 |
| P07632 | ISLSGEHSIIGR | 2,monomethylphosphothione[S]; | 4.2 | 8.4 | 2.000 | 1.46E-02 |
| P02780 | IRGTINSTVTLHDYMK | 0,Diethylphosphate[AnyN-term]; | 8.4 | 17.6 | 2.095 | 8.92E-03 |
| Q9JHB9 | IRGTINSTVTLHDYMK | 0,Diethylphosphate[AnyN-term]; | 8.4 | 17.6 | 2.095 | 8.92E-03 |
| P20059 | CNADPGLSALLSDHR | 0,Pyro-carbamidomethyl[AnyN-termC]; | 5.2 | 11.0 | 2.115 | 9.49E-03 |
| Q63041 | QNNICFDNWWVDEYHTQADHSAAR | 5,Carbamidomethyl[C]; | 5.2 | 11.4 | 2.192 | 3.27E-02 |
| P02780 | QCFLDQTDK | 0,Gln->pyro-Glu[AnyN-termQ];2,Carbamidomethyl[C]; | 139.0 | 305.6 | 2.199 | 1.29E-02 |
| P22282 | TPGETMYYISLPGSVR | 6,Oxidation[M]; | 7.4 | 16.6 | 2.243 | 2.39E-02 |
| P29598 | TDSCSGDSGGPLICNIDGR | 4,Carbamidomethyl[C];14,Carbamidomethyl[C]; | 5.4 | 12.8 | 2.370 | 1.26E-02 |
| P02781 | ELEEFDAPPEAVEANLK | 6,Asp->Lys[D]; | 17.8 | 43.0 | 2.416 | 3.37E-03 |
| P02761 | VFMQHIDVLENSLGFK | 0,Delta_H(2)C(2)[AnyN-term]; | 7.2 | 17.6 | 2.444 | 1.65E-02 |
| P02761 | CEAHGITR | 1,Glutathione[C]; | 11.6 | 28.4 | 2.448 | 1.46E-02 |
| P02780 | ASGSGCSILDEVIR | 6,SulfurDioxide[C]; | 5.0 | 12.4 | 2.480 | 2.13E-02 |
| Q9JHB9 | ASGSGCSILDEVIR | 6,SulfurDioxide[C]; | 5.0 | 12.4 | 2.480 | 2.13E-02 |
| P20767 | NSFTCQVTHEGNTVEK | 1,Deamidated[N];5,Carbamidomethyl[C]; | 4.0 | 10.2 | 2.550 | 5.10E-03 |
| P22282 | LDNCPFEEQTEQLK | 4,Carbamidomethyl[C]; | 66.8 | 177.2 | 2.653 | 5.66E-03 |
| P00758 | HLSQPADITDGVK | 4,Gln->Phe[Q]; | 0.6 | 1.6 | 2.667 | 3.41E-02 |
| P07647 | LVSQSFQHPDYIPVFMR | 16,Oxidation[M]; | 1.8 | 4.8 | 2.667 | 2.85E-02 |
| P36373 | HLSQPADITDGVK | 4,Gln->Phe[Q]; | 0.6 | 1.6 | 2.667 | 3.41E-02 |
| P01835 | DSTYSMSSTLSLTK | 6,Carbamidomethyl[M]; | 2.2 | 6.2 | 2.818 | 3.09E-02 |
| P01836 | DSTYSMSSTLSLTK | 6,Carbamidomethyl[M]; | 2.2 | 6.2 | 2.818 | 3.09E-02 |
| P22057 | QGHDTVQPNFQQDK | 0,Gln->pyro-Glu[AnyN-termQ]; | 1.2 | 3.4 | 2.833 | 4.18E-03 |
| P98158 | GRTCKVTGSENPLLVVASR | 0,3sulfo[AnyN-term]; | 1.2 | 3.4 | 2.833 | 2.95E-02 |
| P07647 | AYDHNNDLMLLHLSK | 6,Deamidated[N]; | 6.2 | 17.6 | 2.839 | 3.98E-03 |
| P22282 | LDNCPFEEQTEQLKR | 4,Carbamidomethyl[C]; | 70.4 | 205.6 | 2.920 | 4.71E-02 |
| P02761 | VFMQHIDVLENSLGFK | 12,Formyl[S](Ser->Asp[S]); | 5.8 | 18.4 | 3.172 | 1.99E-02 |
| P06760 | YRQPLRESGPTLDMPVPSSFNDITQEAELR | 7,Cation_Ni[II][E]; | 0.8 | 2.6 | 3.250 | 3.67E-02 |
| P22282 | TPGETMYYISLPGSVR | 4,Cation_Na[E]; | 23.6 | 77.0 | 3.263 | 6.76E-03 |
| P02761 | ETFQLMVLYGR | 0,Diisopropylphosphate[AnyN-term]; | 1.4 | 4.8 | 3.429 | 4.34E-02 |
| P42854 | SSGNSGQNVWIGLHDPTLGQEPNR | 4,Deamidated_18O(1)[N](Delta_H(1)N(-1)18O(1)[N]); | 7.0 | 24.4 | 3.486 | 1.91E-02 |
| P01835 | VLDSVTDQDSK | 0,mTRAQ_13C(6)15N(2)[AnyN-term]; | 1.6 | 5.8 | 3.625 | 2.77E-02 |
| P01836 | VLDSVTDQDSK | 0,mTRAQ_13C(6)15N(2)[AnyN-term]; | 1.6 | 5.8 | 3.625 | 2.77E-02 |
| P29598 | KPSSTVDQQGFQCGQK | 13,Carbamidomethyl[C]; | 3.8 | 14.8 | 3.895 | 3.07E-02 |
| P02782 | EMYNAPPAAVEAK | 0,Unknown_250[AnyN-term]; | 2.2 | 8.8 | 4.000 | 1.00E-02 |
| P14562 | CNSEEHIFVSK | 1,Carbamidomethyl[C]; | 0.4 | 1.6 | 4.000 | 3.88E-03 |
| P36373 | NLLEDEPFAQHR | 0,dNIC[AnyN-term]; | 1.8 | 7.6 | 4.222 | 6.48E-03 |
| P36374 | NLLEDEPFAQHR | 0,dNIC[AnyN-term]; | 1.8 | 7.6 | 4.222 | 6.48E-03 |
| P98158 | SSDSFSAASVIFSNGR | 0,C+12[AnyN-term]; | 0.6 | 2.6 | 4.333 | 3.41E-02 |
| P02761 | GETFQLMVLYGR | 0,C+12[AnyN-term]; | 1.2 | 5.4 | 4.500 | 1.71E-02 |
| P02761 | VFMQHIDVLENSLGFK | 7,Propargylamine[D]; | 9.0 | 41.4 | 4.600 | 3.49E-02 |
| P02781 | ELEEFDAPPEAVEANLK | 0,Methyl[AnyN-term]; | 0.6 | 2.8 | 4.667 | 4.02E-02 |
| P60711 | EHPVLLTEAPLNPK | 7,Hex(2)Sulf(1)[T]; | 0.6 | 2.8 | 4.667 | 1.09E-02 |
| P63259 | EHPVLLTEAPLNPK | 7,Hex(2)Sulf(1)[T]; | 0.6 | 2.8 | 4.667 | 1.09E-02 |
| P15684 | MLSSFLTEDLFK | 1,Met->Aha[M]; | 0.2 | 1.0 | 5.000 | 1.61E-02 |
| P02761 | MQHIDVLENSLGFK | 0,SPITC_13C(6)[AnyN-term]; | 2.4 | 14.4 | 6.000 | 3.88E-03 |
| P07647 | AYDHNNDLMLLHLSK | 5,Deamidated_18O(1)[N](Delta_H(1)N(-1)18O(1)[N]); | 1.4 | 8.4 | 6.000 | 3.73E-02 |
| P30919 | PSYQAVEYMR | 1,Pro->Cys[P]; | 0.4 | 2.6 | 6.500 | 2.95E-02 |
| Q4FZV0 | FQSAVQYAECQSK | 10,Carbamidomethyl[C]; | 1.0 | 6.8 | 6.800 | 8.42E-06 |
| P01048 | GTKKDGAETLYSFK | 3,Phosphoadenosine[K]; | 0.2 | 1.4 | 7.000 | 3.27E-02 |
| P02761 | LNGDWFSIVVASNK | 12,Formyl[S](Ser->Asp[S]); | 0.6 | 4.2 | 7.000 | 3.27E-02 |
| P08932 | GTKKDGAETLYSFK | 3,Phosphoadenosine[K]; | 0.2 | 1.4 | 7.000 | 3.27E-02 |
| P32038 | LCDVAGWGVVTHAGR | 2,Nmethylmaleimide[C]; | 0.2 | 1.4 | 7.000 | 3.27E-02 |
| P42854 | IGLHDPTLGQEPNR | 4,His->Xle[H]; | 0.4 | 2.8 | 7.000 | 4.18E-02 |
| P22282 | PGETMYYISLPGSVR | 0,Phenylisocyanate_2H(5)[AnyN-term]; | 2.0 | 14.4 | 7.200 | 4.58E-02 |
| P82450 | TFEISCCSDHQCK | 6,Carbamidomethyl[C];7,Carbamidomethyl[C];12,Carbamidomethyl[C]; | 0.2 | 1.6 | 8.000 | 2.49E-02 |
| P07647 | AYDHNNDLMLLHLSK | 5,Deamidated[N]; | 2.4 | 19.6 | 8.167 | 2.39E-02 |
| P22282 | EICYFVVYPDYIEQNIHAVR | 0,Amidine[AnyN-term]; | 0.2 | 1.8 | 9.000 | 3.49E-02 |
| P81828 | QCFRCTSFDSTGFCHVGR | 5,Carbamidomethyl[C];14,Carbamidomethyl[C]; | 1.0 | 9.4 | 9.400 | 9.64E-03 |
| Q64230 | PVENRQAIMTILDQEPDAR | 0,SPITC_13C(6)[AnyN-term]; | 1.0 | 10.2 | 10.200 | 2.86E-02 |
| P02770 | PPACYGTVLAEFQPLVEEPK | 4,Carbamidomethyl[C]; | 0.2 | 2.2 | 11.000 | 4.74E-02 |
| P22282 | LDNCPFEEQTEQLKR | 3,Deamidated[N];4,Carbamidomethyl[C]; | 0.6 | 7.2 | 12.000 | 3.65E-02 |
| Q63041 | MGASEVAQEVEVR | 0,CLIP_TRAQ_2[AnyN-term]; | 0.4 | 5.6 | 14.000 | 2.89E-03 |
| P07151 | PNIEMSDLSFSK | 2,Tris[N]; | 0.2 | 3.0 | 15.000 | 8.64E-03 |
| P02650 | NTMLGQSTEELR | 2,Pent(1)HexNAc(1)[T]; | 0.2 | 3.2 | 16.000 | 9.01E-03 |
| P22282 | EICYFVVYPDYIEQNIHAVR | 5,Phe->Trp[F]; | 0.2 | 4.4 | 22.000 | 2.22E-02 |
| P36375 | NNLFEDEPFAQYR | 0,Dimethyl_2H(4)[AnyN-term]; | 0.2 | 5.2 | 26.000 | 4.47E-02 |

Appendix Table 9 Differential Modifications in the Urinary Proteome of the Hydrogenated Vegetable Oil Group and the Control Group Screened with FC≥1.5 or ≤0.67, P < 0.05

| Uniprot ID | Peptide | Modification | A组 | E组 | Flod change | P value |
| --- | --- | --- | --- | --- | --- | --- |
| P02761 | IDVLENSLGFK | 1,Dioxidation[I]; | 1.0 | 0.0 | 0.000 | 3.41E-02 |
| P04937 | TNIGPDTMRVTWAPPPSIELTNLLVR | 9,Arg->His[R]; | 1.0 | 0.0 | 0.000 | 3.41E-02 |
| P36373 | NNLLEDEPFAQHR | 3,Xle->Gln[L]; | 1.4 | 0.0 | 0.000 | 2.49E-02 |
| P36374 | NNLLEDEPFAQHR | 3,Xle->Gln[L]; | 1.4 | 0.0 | 0.000 | 2.49E-02 |
| P60711 | MCKAGFAGDDAPR | 2,Nmethylmaleimide[C]; | 0.8 | 0.0 | 0.000 | 1.61E-02 |
| P63259 | MCKAGFAGDDAPR | 2,Nmethylmaleimide[C]; | 0.8 | 0.0 | 0.000 | 1.61E-02 |
| Q63621 | CDDWGLDTMR | 1,Carbamidomethyl[C]; | 3.8 | 0.0 | 0.000 | 3.04E-02 |
| Q9JJ19 | IVEVNGVCMEGK | 8,Carbamidomethyl[C];9,Oxidation[M]; | 4.8 | 0.0 | 0.000 | 3.05E-02 |
| P97574 | MIAEVQEDCYSK | 9,Carbamidomethyl[C]; | 9.0 | 0.0 | 0.000 | 4.59E-02 |
| P14046 | YMVLVPSQLYTETPEK | 2,Oxidation[M]; | 9.6 | 0.4 | 0.042 | 4.59E-02 |
| Q03626 | YMVLVPSQLYTETPEK | 2,Oxidation[M]; | 9.6 | 0.4 | 0.042 | 4.59E-02 |
| Q6IE52 | YMVLVPSQLYTETPEK | 2,Oxidation[M]; | 9.6 | 0.4 | 0.042 | 4.59E-02 |
| P27590 | AHWSDHCCLWSTEIQVK | 4,Hex(1)NeuGc(1)[S]; | 3.8 | 0.2 | 0.053 | 4.91E-02 |
| P02761 | EKIEENGSMRVFMQHIDVLENSLGFK | 0,GIST-Quat[AnyN-term]; | 2.4 | 0.2 | 0.083 | 4.02E-02 |
| P26644 | AVFGCHETYK | 5,Carbamidomethyl[C]; | 14.6 | 1.6 | 0.110 | 3.14E-02 |
| P08426 | IIRHPSYNANTFDNDIMLIK | 7,Tyr->Ala[Y]; | 6.6 | 0.8 | 0.121 | 1.66E-02 |
| Q9JJ19 | IVEVNGVCMEGK | 8,Carbamidomethyl[C]; | 6.6 | 0.8 | 0.121 | 3.37E-02 |
| P02770 | LPCVEDYLSAILNR | 3,CarbamidomethylDTT[C]; | 1.6 | 0.2 | 0.125 | 2.49E-02 |
| P01048 | LESGNQFVLYR | 3,Hex(1)NeuGc(1)[S]; | 4.8 | 0.6 | 0.125 | 4.24E-02 |
| P01048 | DGAETLYSFK | 1,Cation_Li[D]; | 5.0 | 1.0 | 0.200 | 3.74E-02 |
| P08932 | DGAETLYSFK | 1,Cation_Li[D]; | 5.0 | 1.0 | 0.200 | 3.74E-02 |
| P26644 | ITCPPPPIPK | 3,Carbamidomethyl[C]; | 27.8 | 5.8 | 0.209 | 1.17E-03 |
| P83121 | DEICAWVVVTTR | 4,Carbamidomethyl[C]; | 1.8 | 0.4 | 0.222 | 2.49E-02 |
| P81828 | DEICAWVVVTTR | 4,Carbamidomethyl[C]; | 1.8 | 0.4 | 0.222 | 2.49E-02 |
| P81827 | DEICAWVVVTTR | 4,Carbamidomethyl[C]; | 1.8 | 0.4 | 0.222 | 2.49E-02 |
| P24090 | IAATDCTGQEVTDPAK | 0,Galactosyl[AnyN-term]; | 3.0 | 0.8 | 0.267 | 4.02E-02 |
| P07632 | ASGEPVVVSGQITGLTEGEHGFHVHQYGDNTQGCTTAGPHFNPH | 34,Carbamidomethyl[C]; | 4.6 | 1.4 | 0.304 | 2.99E-02 |
| P20059 | GECQSEGVLFFQGNR | 0,Acetyl_2H(3)[AnyN-term]; | 9.6 | 3.6 | 0.375 | 2.17E-02 |
| P02780 | ASGSGCSILDEVIR | 6,SulfurDioxide[C]; | 5.0 | 2.2 | 0.440 | 2.49E-02 |
| Q9JHB9 | ASGSGCSILDEVIR | 6,SulfurDioxide[C]; | 5.0 | 2.2 | 0.440 | 2.49E-02 |
| Q62740 | GYSVPTAACR | 9,Carbamidomethyl[C]; | 16.2 | 7.2 | 0.444 | 1.27E-02 |
| Q64240 | AFAELWAFDAAQGK | 9,Cation_Na[D]; | 2.6 | 1.2 | 0.462 | 2.49E-02 |
| P36373 | NNLLEDEPFAQHR | 1,Ammonia-loss[N]; | 2.0 | 1.0 | 0.500 | 3.41E-02 |
| P36374 | NNLLEDEPFAQHR | 1,Ammonia-loss[N]; | 2.0 | 1.0 | 0.500 | 3.41E-02 |
| Q01177 | CEGETDFICR | 1,Carbamidomethyl[C];9,Carbamidomethyl[C]; | 14.6 | 7.8 | 0.534 | 1.05E-03 |
| P36953 | TINPTVDHCCR | 9,Carbamidomethyl[C];10,Carbamidomethyl[C]; | 15.4 | 8.4 | 0.545 | 4.60E-02 |
| P02761 | EKIEENGSMR | 6,Deamidated[N]; | 22.0 | 12.2 | 0.555 | 2.69E-02 |
| Q62740 | ESGDPSTCAFQR | 8,Carbamidomethyl[C]; | 19.8 | 13.0 | 0.657 | 3.10E-02 |
| P07632 | ISLSGEHSIIGR | 2,monomethylphosphothione[S]; | 4.2 | 9.2 | 2.190 | 4.47E-02 |
| P01836 | VLDSVTDQDSK | 0,mTRAQ_13C(6)15N(2)[AnyN-term]; | 1.6 | 4.2 | 2.625 | 4.86E-02 |
| P01835 | VLDSVTDQDSK | 0,mTRAQ_13C(6)15N(2)[AnyN-term]; | 1.6 | 4.2 | 2.625 | 4.86E-02 |
| P42854 | SSGNSGQNVWIGLHDPTLGQEPNR | 4,Deamidated_18O(1)[N](Delta_H(1)N(-1)18O(1)[N]); | 7.0 | 20.8 | 2.971 | 3.19E-02 |
| P02761 | MQHIDVLENSLGFK | 0,SPITC_13C(6)[AnyN-term]; | 2.4 | 9.0 | 3.750 | 2.27E-02 |
| P08426 | LGEHNIDVVEGGEQFIDAAKIIRHPSYNANTFDNDIMLIK | 20,Carbamyl[K]; | 0.4 | 2.4 | 6.000 | 4.74E-02 |
| P00758 | NLYEDEPFAQHR | 3,monomethylphosphothione[Y]; | 2.8 | 17.0 | 6.071 | 1.52E-02 |
| P01836 | RDGVLDSVTDQDSK | 7,Formyl[S](Ser->Asp[S]); | 0.6 | 3.8 | 6.333 | 2.99E-02 |
| P01835 | RDGVLDSVTDQDSK | 7,Formyl[S](Ser->Asp[S]); | 0.6 | 3.8 | 6.333 | 2.99E-02 |
| P22282 | PGETMYYISLPGSVR | 0,Phenylisocyanate_2H(5)[AnyN-term]; | 2.0 | 13.0 | 6.500 | 3.20E-02 |
| P00762 | LGEHNINVLEGDEQFINAA | 7,Deamidated[N]; | 0.2 | 1.4 | 7.000 | 3.27E-02 |
| Q9R0T4 | DTGVISVVTSGLDR | 4,Val->Cys[V]; | 0.4 | 3.2 | 8.000 | 3.12E-02 |

Appendix Table 10 Differential Modifications in the Urinary Proteome of the Rapeseed Oil Group and the Control Group Screened with FC≥1.5 or ≤0.67, P < 0.05

| Uniprot ID | Peptide | Modification | A组 | F组 | Flod change | P value |
| --- | --- | --- | --- | --- | --- | --- |
| P02761 | FMQHIDVLENSLGFK | 0,Phenylisocyanate_2H(5)[AnyN-term]; | 3.8 | 0.0 | 0.000 | 3.76E-02 |
| P02761 | AKLNGDWFSIVVASNK | 3,Xle->Ala[L]; | 5.6 | 0.0 | 0.000 | 4.29E-02 |
| P02761 | YVMFHLINFK | 8,Deamidated[N]; | 6.6 | 0.0 | 0.000 | 3.18E-02 |
| P02761 | IEENGSMRVFMQHIDVLENSLGFK | 6,Dehydrated[S]; | 0.8 | 0.0 | 0.000 | 1.61E-02 |
| P02761 | WFSIVVASNK | 0,Hex[AnyN-term]; | 3.4 | 0.0 | 0.000 | 2.99E-02 |
| P02761 | IDVLENSLGFK | 1,Dioxidation[I]; | 1.0 | 0.0 | 0.000 | 3.41E-02 |
| P02770 | LPEAQRLPCVEDYLSAILNR | 9,CarbamidomethylDTT[C]; | 1.2 | 0.0 | 0.000 | 3.27E-02 |
| P02770 | LPCVEDYLSAILNR | 3,CarbamidomethylDTT[C]; | 1.6 | 0.0 | 0.000 | 3.49E-02 |
| P05544 | ITGTKDLYVSQVVHK | 0,Carbamyl[AnyN-term]; | 4.8 | 0.0 | 0.000 | 2.61E-02 |
| P01836 | DSTYSMSSTLSLTK | 1,Cation_Na[D]; | 6.8 | 0.0 | 0.000 | 4.00E-02 |
| P01835 | DSTYSMSSTLSLTK | 1,Cation_Na[D]; | 6.8 | 0.0 | 0.000 | 4.00E-02 |
| P24090 | IAATDCTGQEVTDPAK | 0,Galactosyl[AnyN-term]; | 3.0 | 0.0 | 0.000 | 2.85E-02 |
| P04937 | TNIGPDTMRVTWAPPPSIELTNLLVR | 9,Arg->His[R]; | 1.0 | 0.0 | 0.000 | 3.41E-02 |
| P04937 | TNIGPDTMRVTWAPPPSIELTNLLVR | 2,Deamidated[N]; | 2.0 | 0.0 | 0.000 | 2.17E-02 |
| P0DP29 | DGNGYISAAELR | 3,Deamidated[N]; | 4.2 | 0.0 | 0.000 | 4.54E-02 |
| P0DP30 | DGNGYISAAELR | 3,Deamidated[N]; | 4.2 | 0.0 | 0.000 | 4.54E-02 |
| P0DP31 | DGNGYISAAELR | 3,Deamidated[N]; | 4.2 | 0.0 | 0.000 | 4.54E-02 |
| P14046 | KQSGVKEEHSFTVMEFVLPR | 0,SPITC_13C(6)[AnyN-term]; | 2.0 | 0.0 | 0.000 | 4.74E-02 |
| Q03626 | KQSGVKEEHSFTVMEFVLPR | 0,SPITC_13C(6)[AnyN-term]; | 2.0 | 0.0 | 0.000 | 4.74E-02 |
| P05545 | QTQGKIAELFSELDER | 0,GIST-Quat_2H(3)[AnyN-term]; | 3.8 | 0.0 | 0.000 | 3.76E-02 |
| P32038 | DVAGWGVVTHAGR | 0,mTRAQ_13C(6)15N(2)[AnyN-term]; | 2.2 | 0.0 | 0.000 | 2.95E-02 |
| P32038 | QQLTVSIMDR | 0,ISD_z+2_ion[AnyN-term]; | 0.8 | 0.0 | 0.000 | 1.61E-02 |
| Q6IE52 | KQSGVKEEHSFTVMEFVLPR | 0,SPITC_13C(6)[AnyN-term]; | 2.0 | 0.0 | 0.000 | 4.74E-02 |
| P08723 | LCLTSLSFTTNK | 1,Xle->Tyr[L]; | 1.2 | 0.0 | 0.000 | 3.27E-02 |
| P00758 | QPGDDYSNDLMLLHLSQPADITDGVK | 4,Cation_Na[D]; | 1.8 | 0.0 | 0.000 | 3.67E-02 |
| P00758 | NLYEDEPFAQHR | 0,GIST-Quat[AnyN-term]; | 5.8 | 0.0 | 0.000 | 3.68E-02 |
| P27590 | HTEDTTIQVTENGESSQAR | 1,Cresylphosphate[H]; | 10.4 | 0.0 | 0.000 | 4.51E-02 |
| P27590 | STDYGAGYSCDSDMHGWYR | 10,Propionamide[C]; | 2.6 | 0.0 | 0.000 | 4.86E-02 |
| P27590 | VCQDPCNVYETLTEYWR | 6,4-ONE[C]; | 2.4 | 0.0 | 0.000 | 4.18E-02 |
| P22282 | YIEQNIHAVR | 0,ISD_z+2_ion[AnyN-term]; | 2.0 | 0.0 | 0.000 | 4.74E-02 |
| P07647 | VGSICLASGWGMTNPSEMK | 5,Carbamidomethyl[C]; | 3.4 | 0.0 | 0.000 | 2.99E-02 |
| P07632 | AVCVLKGDGPVQGVIHFEQK | 3,NEMsulfur[C]; | 2.2 | 0.0 | 0.000 | 2.95E-02 |
| P36373 | NNLLEDEPFAQHR | 3,Xle->Gln[L]; | 1.4 | 0.0 | 0.000 | 2.49E-02 |
| P02780 | SGSGCSILDEVIR | 5,monomethylphosphothione[C]; | 8.0 | 0.0 | 0.000 | 4.91E-02 |
| P02780 | ASGSGCSILDEVIR | 0,Acetyl[ProteinN-term]; | 2.4 | 0.0 | 0.000 | 2.40E-02 |
| Q9JHB9 | SGSGCSILDEVIR | 5,monomethylphosphothione[C]; | 8.0 | 0.0 | 0.000 | 4.91E-02 |
| Q9JHB9 | ASGSGCSILDEVIR | 0,Acetyl[ProteinN-term]; | 2.4 | 0.0 | 0.000 | 2.40E-02 |
| P01048 | QEEGAQELNCNDETVFQAVDTALK | 0,Gln->pyro-Glu[AnyN-termQ];10,Carbamidomethyl[C]; | 3.8 | 0.0 | 0.000 | 2.36E-02 |
| P01048 | LESGNQFVLYR | 3,Hex(1)NeuGc(1)[S]; | 4.8 | 0.0 | 0.000 | 3.49E-02 |
| P36374 | NNLLEDEPFAQHR | 3,Xle->Gln[L]; | 1.4 | 0.0 | 0.000 | 2.49E-02 |
| P06866 | IEDDSCPKPPEIANGYVEHLVR | 6,NEIAA_2H(5)[C]; | 2.0 | 0.0 | 0.000 | 3.41E-02 |
| P31044 | GNDISSGTVLSEYVGSGPPK | 3,Cation_Na[D]; | 4.6 | 0.0 | 0.000 | 4.02E-02 |
| Q68FP1 | TPSAAYLWVGTGASDAEK | 2,Delta_H(5)C(2)[P]; | 2.0 | 0.0 | 0.000 | 4.74E-02 |
| P09005 | ITGTKDLYVSQVVHK | 0,Carbamyl[AnyN-term]; | 4.8 | 0.0 | 0.000 | 2.61E-02 |
| Q63083 | LVTLEEFLASTQR | 3,Dehydrated[T]; | 1.0 | 0.0 | 0.000 | 3.41E-02 |
| P22283 | LVNCPFEEQTEQLK | 4,dichlorination[C]; | 1.4 | 0.0 | 0.000 | 2.49E-02 |
| Q63621 | CDDWGLDTMR | 1,Carbamidomethyl[C]; | 3.8 | 0.0 | 0.000 | 3.04E-02 |
| P19218 | CLLGGLGFK | 1,Carbamidomethyl[C]; | 10.2 | 0.0 | 0.000 | 3.93E-02 |
| P12788 | HPEYNKDTLDNDIMLIK | 0,TMT2plex[AnyN-term]; | 2.0 | 0.0 | 0.000 | 2.17E-02 |
| Q99PS8 | HPPVFGLCR | 8,Carbamidomethyl[C]; | 9.0 | 0.0 | 0.000 | 3.67E-02 |
| Q9JJ19 | IVEVNGVCMEGK | 8,Carbamidomethyl[C]; | 6.6 | 0.0 | 0.000 | 1.75E-02 |
| Q9JJ19 | IVEVNGVCMEGK | 8,Carbamidomethyl[C];9,Oxidation[M]; | 4.8 | 0.0 | 0.000 | 3.05E-02 |
| P20059 | GECQSEGVLFFQGNR | 0,Acetyl_2H(3)[AnyN-term]; | 9.6 | 0.0 | 0.000 | 4.59E-02 |
| P20059 | YYCFQGNK | 3,Carbamidomethyl[C]; | 11.2 | 0.0 | 0.000 | 4.64E-02 |
| P97574 | MIAEVQEDCYSK | 9,Carbamidomethyl[C]; | 9.0 | 0.0 | 0.000 | 4.59E-02 |
| P07314 | DIQEAGGIMTVEDLNNYR | 0,Acetyl_2H(3)[AnyN-term]; | 3.2 | 0.0 | 0.000 | 4.01E-02 |
| P08592 | EVCSEQAETGPCR | 3,Carbamidomethyl[C];12,Carbamidomethyl[C]; | 18.0 | 0.0 | 0.000 | 2.25E-02 |
| P08592 | STNLHDYGMLLPCGIDK | 13,Carbamidomethyl[C]; | 7.0 | 0.0 | 0.000 | 1.92E-02 |
| O70535 | EIICSWNPGR | 4,Carbamidomethyl[C]; | 6.4 | 0.0 | 0.000 | 3.88E-02 |
| P14562 | YSGTCGAQLVTLK | 5,Carbamidomethyl[C]; | 4.0 | 0.0 | 0.000 | 4.74E-02 |
| P14630 | TDLFSISCPGGIMLK | 8,Carbamidomethyl[C]; | 5.2 | 0.0 | 0.000 | 1.69E-02 |
| P80067 | AISYCHETMTGWVHDVLGR | 5,Carbamidomethyl[C]; | 3.4 | 0.0 | 0.000 | 2.99E-02 |
| Q5FVR0 | DEMVPTCWGR | 7,Carbamidomethyl[C]; | 9.6 | 0.0 | 0.000 | 3.55E-02 |
| P02761 | VFMQHIDVLENSLGFK | 4,Gln->Tyr[Q]; | 9.0 | 0.2 | 0.022 | 3.28E-02 |
| P82450 | QCAVYHTSSVLPAPPFTAR | 2,Carbamidomethyl[C]; | 8.8 | 0.2 | 0.023 | 1.87E-02 |
| P02761 | NLDVAKLNGDWFSIVVASNK | 6,Xlink_DTSSP[174][K]; | 16.0 | 0.4 | 0.025 | 4.53E-02 |
| P00787 | HCGIESEIVAGIPR | 2,Cytopiloyne[C]; | 8.0 | 0.2 | 0.025 | 2.47E-02 |
| P02761 | GETFQLMVLYGR | 0,Ethylphosphate[AnyN-term]; | 23.0 | 0.6 | 0.026 | 4.56E-02 |
| P83121 | HICQTYPDEICAWVVVTTR | 3,NEM_2H(5)[C]; | 118.4 | 3.2 | 0.027 | 4.91E-02 |
| P08426 | IIRHPSYNANTFDNDIMLIK | 7,Tyr->Ala[Y]; | 6.6 | 0.2 | 0.030 | 1.83E-02 |
| P10960 | EVVDSYLPVILDMIK | 13,Oxidation[M]; | 5.2 | 0.2 | 0.038 | 2.04E-02 |
| P47853 | VVQCSDLGLK | 4,Carbamidomethyl[C]; | 13.2 | 0.6 | 0.045 | 4.47E-02 |
| P02770 | AMCTSFQENPTSFLGHYLHEVAR | 3,Carbamidomethyl[C]; | 4.2 | 0.2 | 0.048 | 3.74E-02 |
| P83121 | ICQTYPDEICAWVVVTTR | 2,GG[C](Dicarbamidomethyl[C]);10,Carbamidomethyl[C]; | 6.4 | 0.4 | 0.063 | 2.31E-02 |
| P81827 | ICQTYPDEICAWVVVTTR | 2,GG[C](Dicarbamidomethyl[C]);10,Carbamidomethyl[C]; | 6.4 | 0.4 | 0.063 | 2.31E-02 |
| P0DP29 | QLTEEQIAEFK | 3,PhosphoHex[T]; | 6.0 | 0.4 | 0.067 | 3.29E-02 |
| P0DP30 | QLTEEQIAEFK | 3,PhosphoHex[T]; | 6.0 | 0.4 | 0.067 | 3.29E-02 |
| P0DP31 | QLTEEQIAEFK | 3,PhosphoHex[T]; | 6.0 | 0.4 | 0.067 | 3.29E-02 |
| Q5U206 | QLTEEQIAEFK | 3,PhosphoHex[T]; | 6.0 | 0.4 | 0.067 | 3.29E-02 |
| P15083 | SVSCDQSSQIVSMTLNPVK | 0,Unknown_250[AnyN-term]; | 5.6 | 0.4 | 0.071 | 3.87E-02 |
| P27590 | DWMSIVTPAR | 3,Carbamidomethyl[M]; | 11.0 | 0.8 | 0.073 | 2.56E-02 |
| P02761 | VFMQHIDVLENSLGFK | 2,Phe->Trp[F]; | 20.4 | 1.6 | 0.078 | 2.45E-02 |
| P01048 | DGAETLYSFK | 1,Cation_Li[D]; | 5.0 | 0.4 | 0.080 | 1.69E-02 |
| P08932 | DGAETLYSFK | 1,Cation_Li[D]; | 5.0 | 0.4 | 0.080 | 1.69E-02 |
| P02761 | EKIEENGSMRVFMQHIDVLENSLGFK | 0,GIST-Quat[AnyN-term]; | 2.4 | 0.2 | 0.083 | 4.02E-02 |
| P83121 | PDEICAWVVVTTR | 5,Cys->Oxoalanine[C]; | 2.4 | 0.2 | 0.083 | 2.95E-02 |
| P81828 | PDEICAWVVVTTR | 5,Cys->Oxoalanine[C]; | 2.4 | 0.2 | 0.083 | 2.95E-02 |
| P81827 | PDEICAWVVVTTR | 5,Cys->Oxoalanine[C]; | 2.4 | 0.2 | 0.083 | 2.95E-02 |
| P02780 | LVKPYVQDHFTEK | 0,Carbamyl[AnyN-term]; | 13.6 | 1.2 | 0.088 | 4.05E-02 |
| P02761 | EENGSMRVFMQHIDVLENSLGFK | 8,Val->Asn[V]; | 2.2 | 0.2 | 0.091 | 4.74E-02 |
| Q9JHY1 | VYSPQTAVQVPENDSVK | 0,Phenylisocyanate_2H(5)[AnyN-term]; | 8.2 | 0.8 | 0.098 | 3.11E-02 |
| P02761 | LCEAHGITR | 2,Cysteinyl[C]; | 2.0 | 0.2 | 0.100 | 3.67E-02 |
| P24090 | ELACDDPETEHVALIAVDYLNK | 4,Carbamidomethyl[C]; | 12.8 | 1.4 | 0.109 | 2.00E-02 |
| Q68FP1 | AAYLWVGTGASDAEK | 0,LG-anhydrolactam[AnyN-term]; | 3.6 | 0.4 | 0.111 | 4.01E-02 |
| P27590 | CPHTEDTTIQVTENGESSQAR | 2,Pro->Pyrrolidinone[P]; | 44.8 | 5.8 | 0.129 | 2.94E-02 |
| P07632 | ASGEPVVVSGQITGLTEGEHGFHVHQYGDNTQGCTTAGPHFNPH | 34,Carbamidomethyl[C]; | 4.6 | 0.6 | 0.130 | 2.47E-02 |
| P83121 | CQTYPDEICAWVVVTTR | 9,DTT[C]; | 110.2 | 14.6 | 0.132 | 2.79E-02 |
| P81828 | CQTYPDEICAWVVVTTR | 9,DTT[C]; | 110.2 | 14.6 | 0.132 | 2.79E-02 |
| P81827 | CQTYPDEICAWVVVTTR | 9,DTT[C]; | 110.2 | 14.6 | 0.132 | 2.79E-02 |
| P06866 | YVMLPVADQEK | 3,Oxidation[M]; | 11.8 | 1.6 | 0.136 | 3.58E-02 |
| P27590 | FSIQMFR | 5,Oxidation[M]; | 293.2 | 40.0 | 0.136 | 2.85E-02 |
| P02780 | ASGSGCSILDEVIR | 0,Amidine[AnyN-term]; | 1.4 | 0.2 | 0.143 | 3.27E-02 |
| Q9JHB9 | ASGSGCSILDEVIR | 0,Amidine[AnyN-term]; | 1.4 | 0.2 | 0.143 | 3.27E-02 |
| P15083 | SSVTFECDLGR | 7,Carbamidomethyl[C]; | 52.8 | 9.4 | 0.178 | 3.26E-02 |
| P11232 | AFQEALAAAGDK | 0,Pyridylacetyl[AnyN-term](Phenylisocyanate[AnyN-term]); | 2.2 | 0.4 | 0.182 | 3.67E-02 |
| P02770 | TVMGDFAQFVDK | 1,Thr->Val[T]; | 3.2 | 0.8 | 0.250 | 4.18E-02 |
| P27590 | CPHTEDTTIQVTENGESSQAR | 1,Carbamidomethyl[C]; | 629.8 | 162.2 | 0.258 | 4.96E-02 |
| O35568 | DIDECDIVPDACK | 5,Carbamidomethyl[C];12,Carbamidomethyl[C]; | 10.0 | 2.6 | 0.260 | 2.75E-02 |
| P10252 | ILEYFPNGK | 7,Deamidated[N]; | 26.6 | 7.0 | 0.263 | 3.23E-02 |
| P02761 | NGETFQLMVLYGR | 8,Oxidation[M]; | 215.8 | 67.0 | 0.310 | 3.03E-02 |
| P02782 | NAPPAAVEAK | 0,CAF[AnyN-term]; | 2.4 | 0.8 | 0.333 | 3.49E-02 |
| P01836 | KWKIDGSEQRDGVLDSVTDQDSK | 3,Xlink_DTSSP[174][K]; | 1.6 | 0.6 | 0.375 | 3.41E-02 |
| P26644 | ITCPPPPIPK | 3,Carbamidomethyl[C]; | 27.8 | 12.6 | 0.453 | 2.26E-02 |
| P36374 | KPGNDYSNDLMLLHLK | 11,Oxidation[M]; | 5.0 | 14.2 | 2.840 | 4.97E-02 |
| P00762 | TLNNDIMLIK | 3,Deamidated[N]; | 3.2 | 10.2 | 3.188 | 1.07E-02 |
| P00763 | TLNNDIMLIK | 3,Deamidated[N]; | 3.2 | 10.2 | 3.188 | 1.07E-02 |
| Q6IMF3 | DYQELMNTK | 6,Oxidation[M]; | 8.6 | 28.0 | 3.256 | 4.41E-02 |
| Q63041 | VEPGMAPVAK | 5,Oxidation[M]; | 2.2 | 8.4 | 3.818 | 4.23E-02 |
| P27590 | CPHTEDTTIQVTENGESSQAR | 1,Carbamidomethyl[C];14,Deamidated[N]; | 1.0 | 4.8 | 4.800 | 4.50E-02 |
| P00762 | TLNNDIMLIK | 3,Deamidated[N];7,Oxidation[M]; | 5.2 | 25.0 | 4.808 | 8.11E-04 |
| P00763 | TLNNDIMLIK | 3,Deamidated[N];7,Oxidation[M]; | 5.2 | 25.0 | 4.808 | 8.11E-04 |
| P42854 | SSGNSGQNVWIGLHDPTLGQEPNR | 4,Deamidated[N];8,Deamidated[N]; | 1.0 | 5.4 | 5.400 | 2.91E-03 |
| P02770 | TCVADENAENCDK | 2,CarbamidomethylDTT[C];11,Carbamidomethyl[C]; | 1.6 | 11.4 | 7.125 | 4.09E-02 |
| P22282 | LDNCPFEEQTEQLKR | 4,CarbamidomethylDTT[C]; | 2.0 | 14.8 | 7.400 | 2.13E-02 |
| P02770 | APQVSTPTLVEAAR | 8,Formyl[T](Thr->Glu[T]); | 1.6 | 12.2 | 7.625 | 4.08E-02 |
| P02761 | LCEAHGITR | 8,Sulfo[T]; | 1.4 | 10.8 | 7.714 | 1.63E-02 |
| P22282 | EICYFVVYPDYIEQNIHAVR | 0,Amidine[AnyN-term]; | 0.2 | 1.8 | 9.000 | 3.49E-02 |
| P22282 | LDNCPFEEQTEQLKR | 4,Carboxymethyl_13C(2)[C]; | 0.8 | 7.8 | 9.750 | 4.83E-02 |
| Q01177 | TAVTAAGTPCQEWAAQEPHSHR | 7,Gly->Glu[G]; | 0.2 | 2.0 | 10.000 | 3.67E-02 |
| P02770 | YMCENQATISSK | 2,Oxidation[M];3,Carbamidomethyl[C];5,Deamidated[N]; | 0.2 | 2.2 | 11.000 | 4.74E-02 |
| P81827 | CESLDSTGLCR | 1,Carbamidomethyl[C];10,CarbamidomethylDTT[C]; | 4.2 | 48.6 | 11.571 | 3.22E-02 |
| P81827 | CESLDSTGLCR | 1,CarbamidomethylDTT[C];10,Carbamidomethyl[C]; | 2.6 | 41.0 | 15.769 | 3.33E-02 |
| Q6P7A9 | AVPTQCDVTPNSR | 6,Carbamidomethyl[C]; | 0.6 | 9.8 | 16.333 | 3.55E-03 |
| Q01177 | TAAGTPCQEWAAQEPHSHR | 7,BADGE[C]; | 0.8 | 16.0 | 20.000 | 5.84E-03 |
| P02770 | TVMGDFAQFVDK | 3,Met->Lys[M]; | 0.2 | 5.8 | 29.000 | 2.64E-02 |
